# Multimodal AI Decodes Extreme Environment Functional Dark Matter Beyond Homology

**DOI:** 10.1101/2025.11.15.688615

**Authors:** Mengyang Xu, Dantong Wang, Qifan Liu, Hong Jiang, Xiaobin Liu, Yao Li, Di Wang, Hao Dong, Xinkai Yan, Yujing Liu, Anren Xu, Haidong Peng, Yanan Zhang, Hanbo Li, Shengying Li, Jianwei Chen, Xianfen Wu, Yang Wang, Denghui Li, Shanshan Liu, Liang Meng, Yuxiang Li, Chuang Xue, Ling Jiang, Yumeng Zhang, Jiangning Song, Minxiao Wang, Yang Guo, Zengpeng Li, Yue Shen, Xian Fu, Thomas Mock, Yunyun Zhuang, Changhu Xue, Jian Wang, Huanming Yang, Xun Xu, Simon Ming Yuen Lee, Guangyi Fan, Xiangzhao Mao

## Abstract

Functional annotation of proteins from extreme environments represents a major bottleneck for bioresource discovery, as a vast reservoir of functional dark matter defies existing homology-based methods. We demonstrate that environmental pressures impart conserved physicochemical energy signatures that co-determine protein function with sequence and structure. Here we developed ACCESS, a multimodal graph neural network employing hierarchical contrastive learning with a tailored label-sample co-embedding to fuse energy, sequence, and structural information and overcome homology scarcity. ACCESS surpasses state-of-the-art methods including BLASTp and CLEAN in annotating low-identity enzymes. Applied to extreme environmental metagenomics, we constructed a function map of extremophile enzymes to expand the biocatalyst library, pinpointed functionally critical residues to guide rational design, and enabled large-scale, function-based macro-evolutionary analyses. This paradigm transcends the limitations of homology, illuminating protein dark matter and accelerating the exploration of the biosphere’s functional diversity for applications in biotechnology and therapeutic development.

## INTRODUCTION

Proteins are the molecular engines and architects of life, executing the vast majority of processes that define a living cell. From catalyzing metabolic reactions to providing structural integrity and mediating cellular communication, their functional diversity forms the bedrock of biological complexity ^1^. The advent of high-throughput sequencing technologies, particularly metagenomics, has driven an unprecedented expansion in known protein sequences, revealing a biosphere with microbial and genetic diversity far beyond historical expectations. Yet, the current capacity to functionally annotate this deluge of sequences lags behind, exposing a profound and expanding ontological gap ^2^. This collection of uncharacterized sequences is broadly referred to as protein dark matter ^3–5^, typically defined by sequence identity below 0.300 relative to annotated proteins. This frontier obscures our view of metabolic pathways, ecological roles, and evolutionary history, limiting efforts to harness nature’s biotechnological potential, from developing novel therapeutics to engineering robust industrial biocatalysts ^6–9^.

The challenge of annotating this dark matter stems not merely from data scale but also from the fundamental limitations of existing tools ^10^. Experimental characterization of protein function on this scale is laborious and economically unfeasible. For decades, homology-based inference has served as the gold standard for computational function prediction ^11–14^. This paradigm, embodied by algorithms such as BLASTp, operates on the principle that a shared ancestry, detectable through sequence or structural similarity, implies conserved function ^15–17^. While this approach has been remarkably successful for well-studied protein families, its efficacy diminishes as we venture further from the well-lit paths of model organisms. More recent approaches, such as CLEAN, leverage sequence embeddings to extend beyond direct similarity, but their predictive power remains anchored to patterns learned from sequence information ^18^. Consequently, their efficacy declines for highly evolutionarily divergent proteins. The evolutionary chasms separating known proteins from novel sequences discovered in unique ecological niches are often too wide for sequence-based methods to cross, leaving the majority of dark matter functionally opaque ^19^.

This limitation is most acute in extreme environments, where proteins have evolved under unique selective pressures such as extreme temperature, pH, salinity or pressure, thus frequently lacking identifiable homologs ^20,21^. The Mariana Trench Environment and Ecology Research (MEER) project was initiated to explore this frontier ecosystem, seeking to uncover the origin, biodiversity, and adaptive strategies of hadal life ^22^. A cornerstone achievement of MEER is the largest and most comprehensive hadal microbial metagenome dataset to date, which has unveiled a staggering degree of biological novelty ^23–25^. Proteins from these organisms have followed such distinct evolutionary trajectories from their terrestrial or shallow-sea counterparts that their sequences and structures bear little resemblance to any known entries in existing databases. Beyond sequence and structural dark matter (sequence identity < 0.300 or TM-score < 0.500), we further delineate this subset termed “functional dark matter” proteins as a more recalcitrant cohort of proteins characterized by a dual absence of significant sequence identity and appreciable structural similarity to any functionally characterized terrestrial or shallow-sea proteins, representing an untapped reservoir of biocatalysts of the MEER proteome.

To illuminate this functional dark matter, we must look beyond sequence and structure to the underlying physical principles that govern function. The central premise of our work is that the canonical sequence-structure-function model based on Anfinsen’s dogma, while foundational, is incomplete for fully describing proteins shaped by extreme evolutionary forces ^26^. We posit that proteins are governed by conserved networks of stabilizing interactions, such as intramolecular backbone and sidechain hydrogen bond energy, geometric potential energy, intermolecular Lennard-Jones potential energy, Coulombic electrostatic potential energy and asymmetric solvation energy, which manifest as distinct physicochemical energy signatures that co-determine function alongside sequence and structure. The extreme physicochemical conditions of the hadal zone such as ultra-high pressure generate energy effects that exert control over protein folding and function. This environmentally encoded energy modality represents a rich yet largely overlooked source of information in the field of AI-driven protein science ^27–30^.

Harnessing this insight requires a computational approach capable of integrating heterogeneous data modalities. To address this need, we present ACCESS (<u>A</u> <u>C</u>ontrastive <u>C</u>ross-modal <u>E</u>nergy-<u>S</u>tructure-<u>S</u>equence framework for protein function prediction), a multimodal message passing neural network architecture to synergistically integrate sequence information encoded by state-of-the-art (SOTA) protein language model ESM-2 ^31^, structure information predicted via ESMFold ^32^ or AlphaFold3 ^33^, energy information quantified using physics-based Rosetta calculations ^34,35^, and hierarchical functional annotations from the Enzyme Commission (EC) number system ^36,37^. The core innovation of ACCESS addresses three critical challenges in dark matter annotation by employing a hierarchical contrastive learning strategy to improve prediction accuracy, combining geometric vector perceptrons and graph attention to integrate heterogeneous energy and sequence information into structure-derived graphs, and constructing a label-sample latent space guided by the EC label hierarchy and co-occurrence network to specifically target homology-scarce proteins. This integrated design allows ACCESS to comprehensively outperform current SOTA methods including BLASTp, CLEAN and those based on transformers and vanilla graph neural networks, across both general protein annotation and dark matter subsets.

By applying ACCESS to large-scale metagenomic datasets from MEER, we have generated a comprehensive functional map of extremophile enzymes that massively expands the known library of natural biocatalyst templates. Beyond annotation, the model’s inherent interpretability enables the high-precision identification of functionally critical residues, including distal allosteric sites, thereby providing an actionable blueprint for rational protein design. Furthermore, our function-first annotation framework facilitates robust, large-scale macro-evolutionary analyses based on functional profiles rather than sequence phylogeny alone. ACCESS thus establishes a new paradigm that transcends the limitations of homology and extends Anfinsen’s dogma by demonstrating environmental energy effects, illuminating the functional dark matter of the protein universe and accelerating the exploration of the biosphere’s functional diversity for transformative applications in biotechnology and therapeutic development.

## RESULTS

### Landscape of Protein Functional Dark Matter in Extreme Environments

The MEER project’s metagenomic surveys of the Mariana Trench have uncovered a remarkable level of microbial novelty, laying the groundwork for investigating protein functional diversity in hadal ecosystems. Analysis of sediment samples revealed thousands of microbial species, with 89.4% representing taxa not previously reported in public databases (Figure 1A) ^38,39^. This high degree of taxonomic novelty directly translates to functional uncertainty at the protein level, as most proteins encoded by these unreported species lack homologs in well-annotated databases. The Deep-Sea Protein Fold Catalog (DSFC) dataset extracted from MEER and NCBI samples represents a collection of 502 million nonredundant genes and 2,400,556 representative gene clusters (≥ 30 members per cluster at > 0.200 amino-acid identity) sourced from microbes, amphipods, and fish, adapted to the extreme conditions of the hadal zone, serving as an unparalleled resource for exploring how environmental pressures shape protein function ^23–25^.

**Figure 1.**
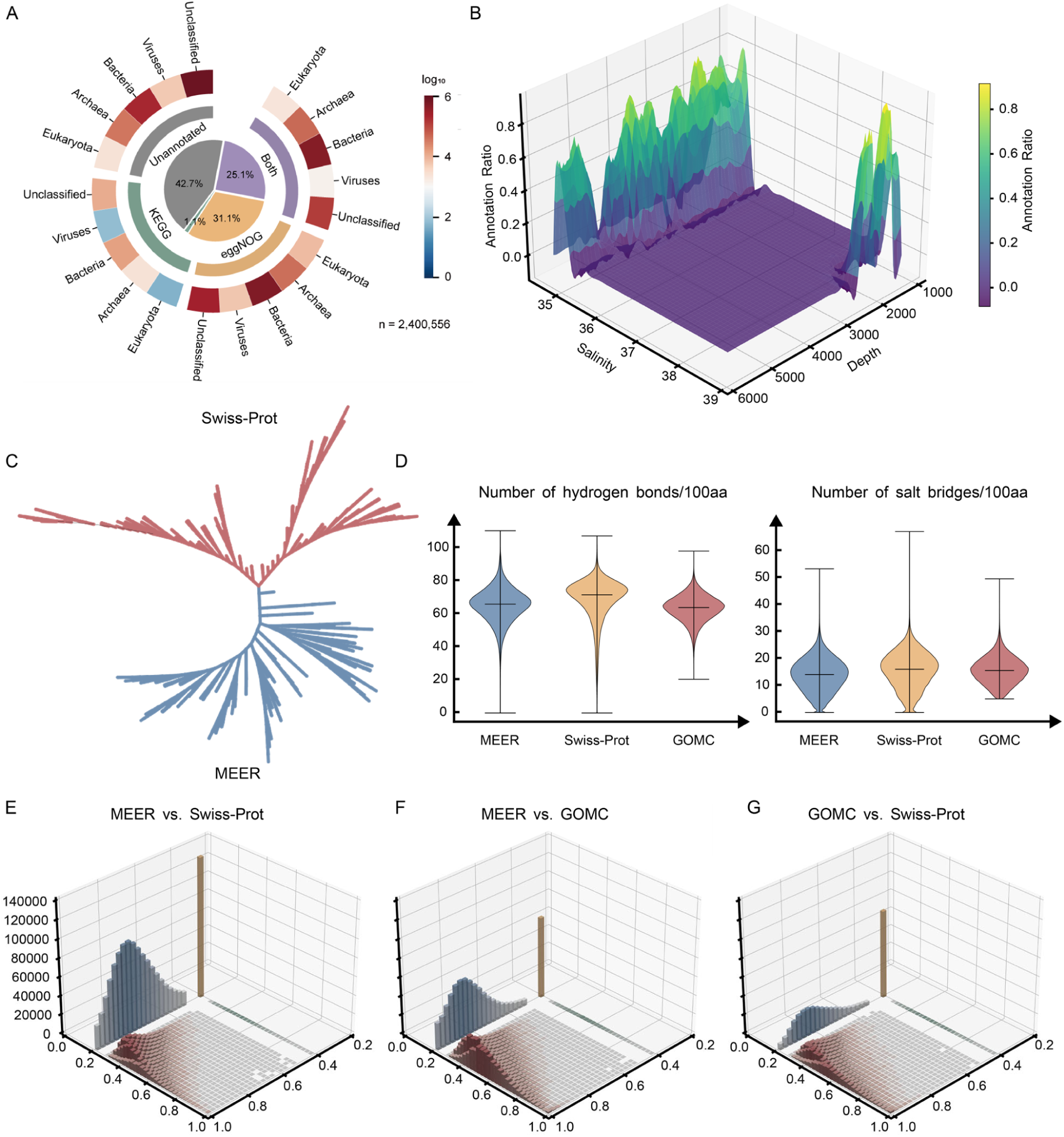
The unique evolutionary and energetic landscape of the Mariana Trench proteome reveals profound limitations of homology-based annotation. (A) A circular proportion chart illustrating the functional annotation bottleneck in the MEER dataset. 42.8% of proteins remain functionally unannotated by SOTA, homology-based tools (eggNOG and KEGG), representing a vast reservoir of functional dark matter. Compounding this functional obscurity further, 48.0% of these unannotated protein sequences originate from species that are taxonomically unclassified below the phylum level, highlighting a parallel taxonomic dark matter that severs ties to known biological contexts. (B) A 3D heatmap revealing the correlation between environmental pressure and the prevalence of unannotated proteins. The axes represent depth (meters below sea level), salinity (Practical Salinity Units, PSU), and the proportion of functionally unannotated proteins from the MEER dataset. The color gradient, from blue (low) to yellow (high), indicates the density of these unannotated proteins, demonstrating that functional dark matter increases systematically with environmental extremity, peaking in the hadal zone. (C) A maximum-likelihood phylogenetic tree contrasting the evolutionary trajectories of orthologous proteins from the deep-sea MEER dataset (blue branches) with their corresponding homologs from the terrestrial- and surface-dominated Swiss-Prot database (red branches). The distinct clustering of MEER orthologs signifies accelerated or divergent evolutionary paths, suggesting neofunctionalization driven by the selective pressures of the deep-sea environment, a process that inherently obscures homologous relationships. (D) Violin plots revealing a statistically significant shift in the energetic properties of deep-sea proteins. Distributions of intramolecular hydrogen bonds and salt bridges per 100 residues are compared across three proteomes: the piezophilic MEER dataset, the mesophilic Swiss-Prot database, and the surface-ocean GOMC catalog. MEER proteins exhibit a statistically higher density of salt bridges (p < 0.001), suggesting an adaptation for stability under high hydrostatic pressure. (E-G) 3D feature-space maps delineating the boundaries of protein novelty. The entire MEER, Swiss-Prot, and GOMC proteomes are projected onto a space defined by sequence similarity (BLASTp E-value) and structural similarity (TM-score). These maps identify distinct uncharacterized zones of protein universe: sequence dark matter (low sequence similarity; red), structural dark matter (low structural similarity; blue), and the functional dark matter (low similarity in both; yellow), a region of extreme novelty where homology-based inference completely fails. The MEER vs. Swiss-Prot (E) comparison shows the largest population of proteins within the functional dark matter, exceeding that of MEER vs. GOMC (F) or GOMC vs. Swiss-Prot (G), empirically confirming the deep-sea biome as a rich source of novel protein folds and functions.

Systematic annotation of 2.4 million representative proteins from the MEER-DSFC gene clusters using conventional sequence homology-based tools revealed that a substantial fraction qualifies as functional dark matter, defined by extreme sequence heterogeneity that precludes reliable matches to experimentally characterized proteins in public databases ^38–40^. To further characterize this dark matter, we analyzed its distribution relative to key environmental parameters in the Mariana Trench. Visualization of the prevalence of unannotated proteins against environmental parameters showed that the highest density of dark matter aligns with zones of extreme pressure and high salinity (Figure 1B). This spatial coincidence suggests that the unique energy blueprint of the abyssal environment acts as a powerful selective force, driving the evolution of novel protein functions and structures that differ from those found in more temperate biomes ^41^.

To explore the disconnect between sequence similarity and functional relatedness in proteins from extreme environments, we conducted phylogenetic analysis of the photosystem I P700 chlorophyll a apoprotein A2 (EC: 1.97.1.12), an enzyme with conserved catalytic function across diverse ecosystems. After identifying proteins from both the MEER and Swiss-Prot databases that exhibit high structural similarity to the query proteins (AlphaFold Protein Structure Database (AFDB) ID: AF-Q7NFT5-F1-model_v4), we clustered their sequences and found that the resulting tree revealed two distinct, well-supported monophyletic clades: one comprising exclusively MEER sequences and the other exclusively known sequences from the reviewed section of a universal protein knowledgebase UniProt (Swiss-Prot database) ^42,43^. This pronounced divergence observed, even within proteins with identical fundamental catalytic function, underscores the inadequacy of sequence similarity as the sole indicator of functional or evolutionary relatedness across disparate ecosystems (Figures 1C, S1A, S1B).

Given the limitations of sequence and structural homology, we investigated whether energy landscape features could serve as a more reliable functional signature for hadal proteins. Using Rosetta, we calculated key energy terms for proteins from three datasets: the deep-sea MEER dataset, the terrestrial-dominated Swiss-Prot database ^42^, and the surface-ocean-dominated Global Ocean Microbiome (GOMC) catalog ^44^. Statistical analysis revealed significant differences in these energy signatures across the three groups. MEER proteins exhibited higher average hydrogen bond stability and stronger salt bridges compared to Swiss-Prot and GOMC proteins, indicating adaptations likely evolved to counteract the destabilizing effects of ultra-high pressure (Figures 1D, S1C). These differences were consistent across diverse protein families.

To further validate the utility of energy signatures, we compared proteins from the MEER, Swiss-Prot and GOMC databases (Figure S1D) and mapped the sequence similarity and structural similarity onto a three-dimensional (3D) space (Figures 1E, 1F, 1G). Most unannotated MEER proteins in Figure 1A clustered in three distinct regions: sequence dark matter with sequence similarity to known proteins < 0.300, structural dark matter with structural similarity < 0.500, and functional dark matter where both sequence and structural similarity were low while energy signatures diverged from all well-annotated proteins. This clustering showed that energy signatures provide a critical orthogonal dimension for distinguishing functional groups, even when sequence and structure fail. Notably, we observed a phenomenon of energy isomerism, in which proteins with dissimilar sequences and structures but nearly identical energy signatures often shared the same or related functions. These observations demonstrate that the energy characteristics may represent a more fundamental and conserved signature of function that persists even when sequence and structure similarity is undetectable.

### ACCESS: A Multimodal AI Framework for Energy-Aware Function Prediction

Conventional protein function prediction tools rely heavily on sequence or structure homology, rendering them inherently limited in decoding functional dark matter derived from extreme environments ^45^. To address this gap, we developed ACCESS, a cross-modal AI framework that employs energy characteristics, shaped by extreme environmental selective pressures, as a functional blueprint to integrate multi-source biological data (Figure 2A). Its design incorporates three core synergistic innovations: a hierarchical multi-label prediction strategy for EC number classification, a hybrid message passing neural network combining geometric vector perceptrons and graph attention (GVPs-GAT) for multimodal fusion ^46,47^, and a contrastive learning framework that ensures robust latent space formation ^48^. The model ingests three complementary biological signals to form a comprehensive functional representation of proteins: (1) Sequence-based features: high-dimensional contextual embeddings generated by ESM-2, a transformer-based protein language model trained on the UniRef50 database, which encodes evolutionary patterns and subtle sequence motifs ^32^. (2) Structure-based features: experimentally determined 3D structures from existing databases ^49^ or high-confidence predictions by protein structure prediction tools like AlphaFold3 ^33^. These features capture residue-specific backbone dihedral angles (φ/ψ), side-chain geometries, and inter-residue contact features, all of which model intricate atomic interactions underlying functional conformations. (3) Energy features: quantitative physicochemical energy profiles computed from energy minimization protocols or molecular dynamics simulations, encompassing both residue-level terms (e.g., fa_rep for steric clashes, hbond_sc for side-chain hydrogen bonds, and salt bridge density) and protein-level terms (e.g., total folding energy, cavity volume, and net charge) ^34,35^.

**Figure 2.**
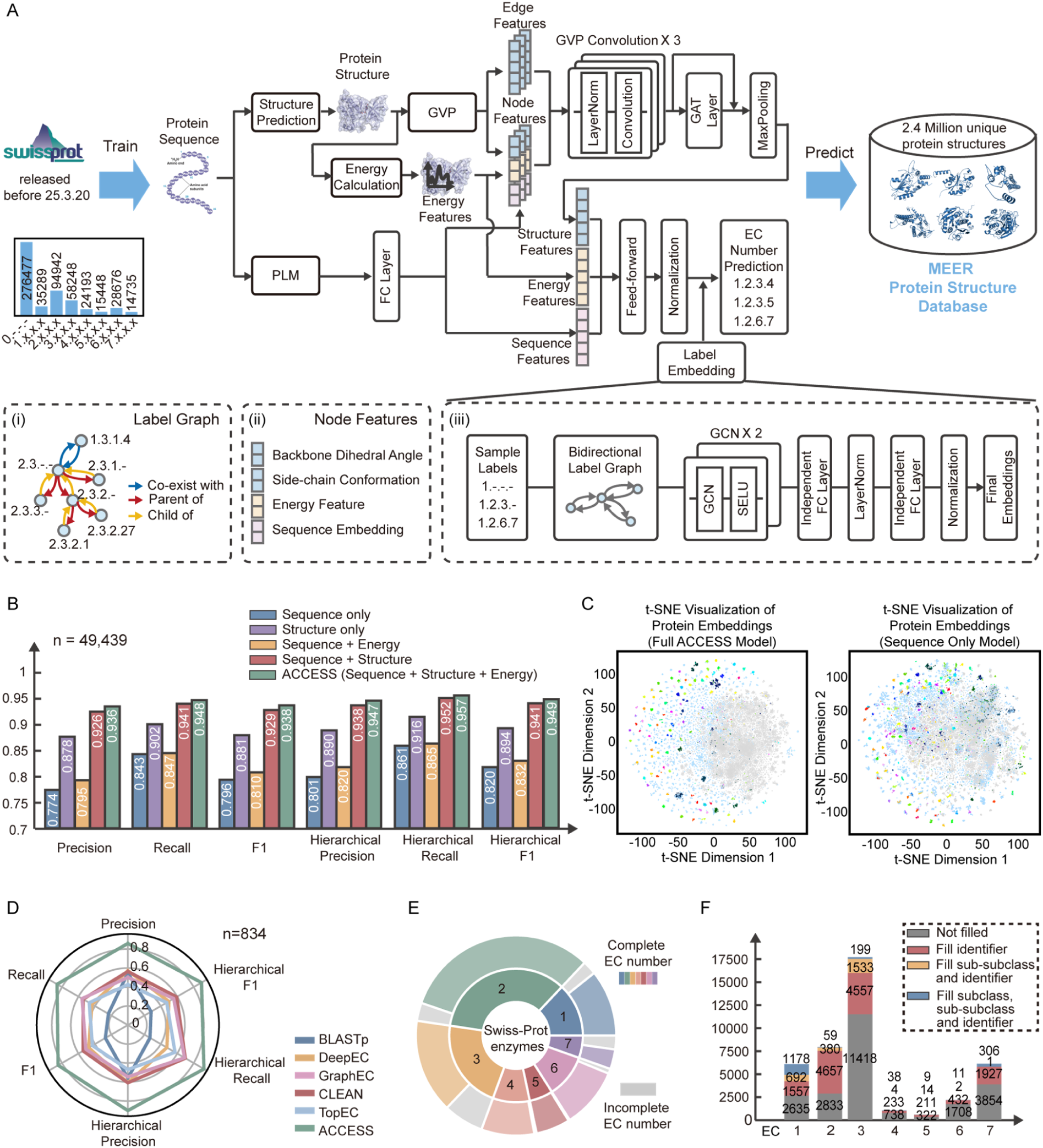
Architecture and benchmark performance of ACCESS, an energy-guided contrastive message-passing neural network for enzyme function prediction. (A) Schematic architecture of ACCESS. The model is trained on 573,661 experimentally validated enzymes from the Swiss-Prot database. It integrates three data modalities as input for a given protein: 1) Sequence: a 1D amino acid sequence processed by a pre-trained protein language model (e.g., extracted from ESM-2) to generate residue-level embeddings; 2) Structure: a 3D coordinate graph of the protein (e.g., derived from AlphaFold3 prediction) processed by geometric vector perceptrons (GVPs) to capture spatial relationships; and 3) Energy: a per-residue energy profile (e.g., calculated using the Rosetta energy function), capturing features like van der Waals forces, solvation energy, and hydrogen bonding potential. These multimodal features are fused as node attributes in a graph representation of the protein, which is then processed by a Message-Passing Neural Network (MPNN). The MPNN iteratively updates residue representations by aggregating information from their spatial neighbors, effectively learning a function-aware representation of the entire protein. A contrastive learning objective function guides the training process, pulling the learned embeddings of proteins towards the vector representations of their true EC number labels while pushing them away from negative labels. In the crucial inference phase, we apply the trained ACCESS model to the MEER dataset with largely unannotated metagenomic data. Three additional insets highlight key innovations: (Inset i) a hierarchical, multi-label loss function: This diagram, using the multi-domain enzyme EC 2.3.2.27 as an example, illustrates how the model’s loss function is designed to respect the four-level EC number hierarchy. It penalizes misclassifications at higher levels of the hierarchy more severely than at lower levels and natively handles proteins with multiple enzymatic functions; (Inset ii) the composition of multimodal node features: backbone dihedral angles and side-chain conformations represent local structure, Rosetta energy features provide biophysical context, and ESM sequence embeddings capture evolutionary information; and (Inset iii) a label-embedding architecture: EC numbers are not treated as simple class indices but are transformed into rich, learnable vector representations. This allows the model to capture semantic relationships between different enzyme functions within the embedding space. (B) Multimodal synergy enhances performance. The bar chart presents the results of a comprehensive ablation study conducted on a hold-out test set of 49,439 proteins from the Swiss-Prot database. Model performance was evaluated using six distinct metrics to provide a robust assessment: standard Precision, Recall, and F1-score, as well as their hierarchical counterparts (H-Precision, H-Recall, H-F1), which account for the tree-like structure of EC numbers. The full ACCESS model, which integrates all three data modalities (’sequence+structure+energy’), achieves higher performance than models trained on any single modality or any combination of two. The results show that each modality provides non-redundant information, with the integration of biophysical energy terms providing performance uplift across all metrics, thereby validating our core hypothesis that the energy landscape is a key determinant of function. (C) Structure and energy features create a discriminative representation space of proteins. This panel displays a t-SNE visualization of protein embeddings, where each point represents a protein and is colored by its EC sub-subclass (e.g., EC 1.1.1). Grey points signify proteins without a formal EC number. The right plot shows the embedding space generated by a model variant trained only on sequence information. While some clustering is evident, class boundaries are diffused. The left plot shows embeddings from the complete ACCESS model, which incorporates sequence, structure, and computationally derived energy features. The incorporation of structure, as well as energy parameters likely calculated from 3D structures to represent thermodynamic stability and interaction potentials, results in dramatically improved separation between functional classes. This demonstrates that structure and energy are functionally relevant modalities that guide the contrastive learning framework to learn a more discriminative and biophysically meaningful representation of protein function. (D) ACCESS outperforms SOTA methods on a general benchmark. The radar chart provides a multi-faceted performance comparison of ACCESS against leading SOTA function prediction tools. The evaluation was conducted on a stringent, non-redundant test set of 834 proteins from Swiss-Prot, which were excluded from the training sets of all benchmarked methods. Performance is assessed across a comprehensive suite of six standard metrics, likely including Precision, Recall, and F1-score, as well as their hierarchical counterparts to ensure a robust evaluation across different aspects of predictive power. (E, F) Large-scale curation and refinement of public protein databases using ACCESS. These panels quantify the practical utility of ACCESS as a high-throughput annotation tool to resolve ambiguity and error in the Swiss-Prot database. The model successfully refined 18,000 proteins with ambiguous, top-level EC annotations (e.g., "EC 3.1.-.-", indicating a generic hydrolase). ACCESS assigned these proteins to specific and complete four-digit EC numbers, significantly increasing the granularity and utility of the functional information available to the scientific community.

ACCESS implements a hierarchical contrastive learning framework whose core innovation lies in jointly mapping both protein samples and their EC labels into a unified embedding space. The protein encoding pathway is built upon a multimodal hybrid message-passing neural network. Each protein is represented as a graph, with nodes corresponding to residues featurized by concatenating sequence, structural, and energetic features, and edges denoting possible inter-residue contacts. GVP layers extract rotation-invariant geometric and structural information from the protein backbone and key side-chain atoms, while the GAT layer captures local structural-energy dependencies. Residue-level outputs are pooled and combined with global sequence and energy features to form a comprehensive protein-level representation, which is then projected via a fully connected layer into a 64D latent space as the final protein embedding. In parallel, a bidirectional graph of EC numbers is constructed to encode the hierarchical and co-occurrence relationships among EC labels. This graph is processed through two graph convolutional network (GCN) layers followed by two independent fully connected layers to project the EC labels into the same latent space as the protein embedding, enabling joint contrastive learning. The model is optimized with a composite loss function designed to align protein and label representations while respecting the EC hierarchy. It combines a joint embedding loss, which minimizes the mean square error (MSE) between a protein’s embedding and that of its ground-truth label, with a hierarchy-aware matching loss. The latter improves discrimination by dynamically identifying the five nearest negative labels for each sample and assigning distinct penalty margins according to their hierarchical relationship to the true label: lightest for parents, moderate for siblings, and strongest for unrelated classes. This design ensures semantic consistency with the EC tree and mitigates misclassifications in sparsely sampled EC subclasses.

During the inference stage, ACCESS assigns one or multiple EC labels to a query protein by adaptively thresholding the Euclidean distances between its embedding and those of the candidate labels, retaining the label(s) whose distance falls within the adaptive margin. For model training and evaluation, we curated a high-quality dataset of 508,645 experimentally validated enzymes from Swiss-Prot. The dataset was randomly split into training, validation, and test sets in an 8:1:1 ratio (406,916, 50,865, and 49,464 proteins, respectively). To explicitly evaluate model generalizability to functional dark matter, defined as proteins with < 0.300 sequence identity and TM-score < 0.500 to any known sequence, we constructed spDarkM-93, a set of 93 Swiss-Prot proteins with no detectable homologs in the training set. This partitioning strategy ensures that ACCESS is not only data-efficient but also specifically validated against the central challenge of extreme environment protein analysis: predicting protein functions in the absence of sequence or structural homologs.

To dissect the contribution of each feature modality and their synergistic integration, we performed systematic ablation studies on the Swiss-Prot test set (n=49,469). We compared five model variants: sequence-only, structure-only, sequence+energy, sequence+structure, and the full ACCESS model (i.e., sequence+structure+energy) in terms of six performance metrics including precision, recall, F1-score, and their hierarchical counterparts ^50^ (Figure 2B). The full ACCESS model consistently outperformed all ablated variants, achieving a precision of 0.936, a recall of 0.948, and an F1-score of 0.938, while the sequence+structure model obtained a precision of 0.926, a recall of 0.941, and an F1-score of 0.929. Notably, removing energy or structural features led to a performance decline in hierarchical F1-scores. These results underscore the complementary roles of each modality: sequence captures evolutionary constraints, structure captures protein folding potential, and energy bridges the gap between folding and function, especially for proteins in extreme environments with decoupled sequence/structure and functional conservation.

A critical evaluation of our multimodal approach is to test its ability to generate a functionally coherent latent space, where proteins with similar functions cluster together and those with different functions are distinctly separated. We employed t-distributed Stochastic Neighbor Embedding (t-SNE) to visualize the 64-dimensional embeddings generated by ACCESS, both with and without the structure and energy features ^51^ (Figure 2C). When these features were excluded, non-enzymatic proteins formed an indistinct cloud (mean nearest-neighbor distance = 0.0693) that overlapped substantially with the cluster representing EC 3 (hydrolases). In contrast, incorporating structure and energy features enabled non-enzymatic proteins to coalesce into a significantly tighter cluster (mean nearest-neighbor distance = 0.0552, paired Wilcoxon test, p *<* 1 x 10^-4^). This enhanced separability confirms that structure and energy features enable ACCESS to excel in two fundamental and challenging classification tasks simultaneously: distinguishing enzymes from non-enzymes and delineating fine-grained functional boundaries among EC numbers. Collectively, these visualizations provide compelling empirical support for our central hypothesis that the physicochemical energy characteristics constitute a critical, yet previously overlooked, dimension for predicting the functions of proteins that lack homologous templates ^52,53^.

We further benchmarked ACCESS’s predictive performance against five SOTA tools, including the sequence-homology-based BLASTp ^17^, the sequence-only deep-learning-based CLEAN ^18^, the sequence-based hierarchy-aware DeepEC ^54^, and the structure-based GraphEC ^55^ and TopEC ^56^. Evaluation was conducted on three datasets: the standard Swiss-Prot test set (n=834) after removing any proteins present in the training or validation splits of the competing methods, a subset of the standard Swiss-Prot test set in which proteins share < 0.300 sequence BLAST identity values with any other protein in our training set, and the purpose-built spDarkM-93 functional dark matter test set (n=93), in which proteins share < 0.300 sequence BLAST identity values and < 0.500 structural TM-scores with any other protein in our training set (Figures 2D, S2A, S2B). On the standard Swiss-Prot test set, ACCESS achieved a hierarchical F1-score of 0.894, outperforming all other methods by 0.301-0.612 (Figure 2D). BLASTp and CLEAN plummeted on this dataset, underscoring the limitations of traditional alignment-based annotation. The structure-based methods, GraphEC and TopEC, while more robust, were still unable to match the performance of ACCESS, suggesting that their reliance on known active site geometries was a constraint when faced with true novelty. The performance gap reemerged on the subset of low-sequence-identity proteins and the spDarkM-93 dark matter test set, where homology-based and conventional deep learning methods are expected to falter (Figures S2A, S2B). For instance, ACCESS correctly assigned EC 4.6.6.1 (Type VI adenylyl cyclase) to a low-sequence-identity pair (Figure S2C) and EC 3.2.2.1 (*α*-D-glucopyranose) to a pair of dark-matter enzymes that share only 0.253 sequence identity and a TM-score of 0.308, whereas all competing methods failed (Figure S2D).

A major bottleneck in protein function prediction is the curse of rarity, where novel or specialized enzyme classes are sparsely sampled in public databases, leading to poor model generalization. ACCESS’s exceptional resilience in these low-data regimes makes it a crucial tool for exploring the unique biochemistry of extremophiles (Figures S2E, S2F). We further focused our analysis on EC Class 7 (translocases), a group enriched in proteins adapted to extreme environments yet notoriously underrepresented in training data, with fewer than 50 training samples available for most of its subclasses. Despite these constraints, ACCESS maintained a hierarchical F1-score of 0.880 (Figure S2F). A paradigmatic case is EC 7.2.2.21 (Cd^2+^-exporting ATPase), an enzyme vital for heavy metal detoxification in specific microbial niches ^57^. With only 16 training samples available and < 0.250 sequence identity to known homologs, ACCESS achieved an F1-score of 1.000. Beyond predictive accuracy, computational scalability is imperative for annotating large-scale metagenomic datasets, where ACCESS outperforms all competitors in both throughput efficiency and resource utilization. When processing the high-volume dataset of 2.4 million protein samples, most existing SOTA methods fail to complete the task under reasonable resource and time constraints. In contrast, ACCESS circumvents computationally expensive preprocessing and alignment steps, and successfully processes the entire dataset within 15 hours (Table 1)

**Table 1.** Comparison of the execution time and peak memory usage of ACCESS, BLASTp, DeepEC, GraphEC, CLEAN, and TopEC.

| Tools | Sample size n=834 |  | Sample size n=2,400,556 |  |
| --- | --- | --- | --- | --- |
|  | Execution time (h) | Peak memory usage (GB) | Execution time (h) | Peak memory usage (GB) |
| ACCESS | 0.0464 | 19.9 | 14.6 | 200 |
| BLASTp | 0.0469 | 0.198 | 5.44 | 12.8 |
| DeepEC | 0.0272 | 2.10 | N/A | N/A |
| GraphEC | 0.159 | 108 | N/A | N/A |
| CLEAN | 0.199 | 16.8 | N/A | N/A |
| TopEC | 0.431 | 7.41 | N/A | N/A |

All tools ran on the same Linux system equipped with an Intel Core Processor (Broadwell, IBRS) of 40 CPU threads and 512 GB RAM. When processing the high-volume dataset of 2.4 million protein samples, DeepEC fails during batch processing due to its excessive memory footprint; CLEAN relies on the CPU-based computation of ESM embeddings as a preprocessing step, making it extraordinarily slow for massive datasets; graph-based TopEC and GraphEC exhibit low inference throughput, as TopEC requires roughly 11 days and GraphEC about 20 days to process 2.4 million samples. BLASTp, though memory-efficient, becomes laboriously infeasible for manual analyses at this scale.

Beyond function prediction, ACCESS enables the correction of ambiguous or inaccurate EC annotations in existing databases (Figures 2E, 2F, S2G). Annotation errors and their propagation within well-established databases are a well-documented challenge in bioinformatics ^58,59^. Across 548,008 Swiss-Prot proteins, ACCESS identified 6.87% as potentially problematic and proposed corresponding corrections falling into three mechanistically distinct categories. The functional activation category involves the assignment of a specific enzymatic function to 12,747 (2.33%) proteins previously labeled as “hypothetical”, “uncharacterized” or “non-enzymatic”. This is achieved by recognizing conserved energy signatures that strongly correlate with known EC classes. A representative example is a hypothetical protein (UniProt ID: Q79G04) that might be assigned by ACCESS as a PET hydrolase (EC 3.1.1.74). The functional reassignment category includes 6,889 proteins (1.26%) with an existing but likely incorrect EC number. For instance, ACCESS flagged a kinase (UniProt ID: Q8IV63) initially labeled EC 2.7.11.22 and reannotated as EC 2.7.11.1 (non-specific serine/threonine kinase). The functional refinement category pertains to 18,000 proteins (3.28%) with vague or incomplete annotations, such as a lipase (UniProt ID: P0A8V0) that ACCESS reassigned from the nonspecific annotation EC 3.1.-.- to the precise function EC 3.1.26.11 (tRNase Z).

### The MEER-EC Functional Atlas Unveils a Reservoir of Enzyme Innovation in Deep Sea

To validate the scalability and utility of our framework, we deployed ACCESS to functionally annotate the 2.4 million non-redundant protein sequences from the MEER project’s metagenomic dataset. This effort culminated in the creation of the MEER-EC Atlas, a comprehensive resource of predicted enzyme functions from one of the planet’s most extreme marine environments. This atlas not only expands the landscape of known enzymatic diversity but also provides a unique perspective on how extreme hadal conditions, particularly hydrostatic pressure and salinity gradients, drive molecular adaptation and catalyze functional innovation in microbial enzymes ^20,41,60,61^.

Our analysis demonstrates that ACCESS penetrates the veil of functional dark matter. The model assigned EC numbers to 612,310 previously unannotated proteins, thereby illuminating a vast reservoir of previously cryptic biochemical potential. Geographic visualization of the MEER-EC Atlas unveils a global biogeography of enzymatic novelty, revealing that these newly annotated functions are widely distributed, mirroring the ubiquity of extreme environments from Arctic cold seeps to the deepest hadal zones (Figures 3A, 3B, S3A, S3B). Examples include novel chitinases (EC 3.2.1.14) crucial for chitin degradation ^62^ identified in Arctic cold seeps samples, and diverse PET-degrading enzymes (EC 3.1.1.74) detected in the abyssal plains of the Mariana Trench ^63^. A notable feature of the atlas is the influence of the bathyal gradient on enzymatic prevalence. Nearly half of the newly annotated enzymes originate from abyssal and hadal environments (depths > 5000 meters) (Figures 3C, S3C), and 7.90% of these show no sequence or structural similarity to any known Swiss-Prot entries (Figure 3D). This underscores the limitations of conventional homology-based annotation that relies on similarity to well-characterized proteins often sourced from less extreme environments. The newly identified enzymes derive from bacteria, archaea, eukaryotes, and viruses, and span all seven major EC classes, revealing the diverse metabolic capabilities and versatility of microbial communities thriving under hadopelagic conditions (Figure 3E). Transferases (EC 2) constitute the most abundant class (36.8%), followed by hydrolases (EC 3, 30.1%) and oxidoreductases (EC 1, 12.7%); lyases, isomerases, ligases, and translocases together comprise the remaining 20.4%. This distribution contrasts with that of other microbial ecosystems and databases, where oxidoreductases (EC 1) often dominate ^64^. To facilitate the research community’s access, the MEER-EC Atlas is hosted on an interactive online portal that allows querying by EC number, specific energy features, or habitat of origin, enabling researchers to prioritize enzymes for experimental validation based on both function and environmental adaptation.

**Figure 3.**
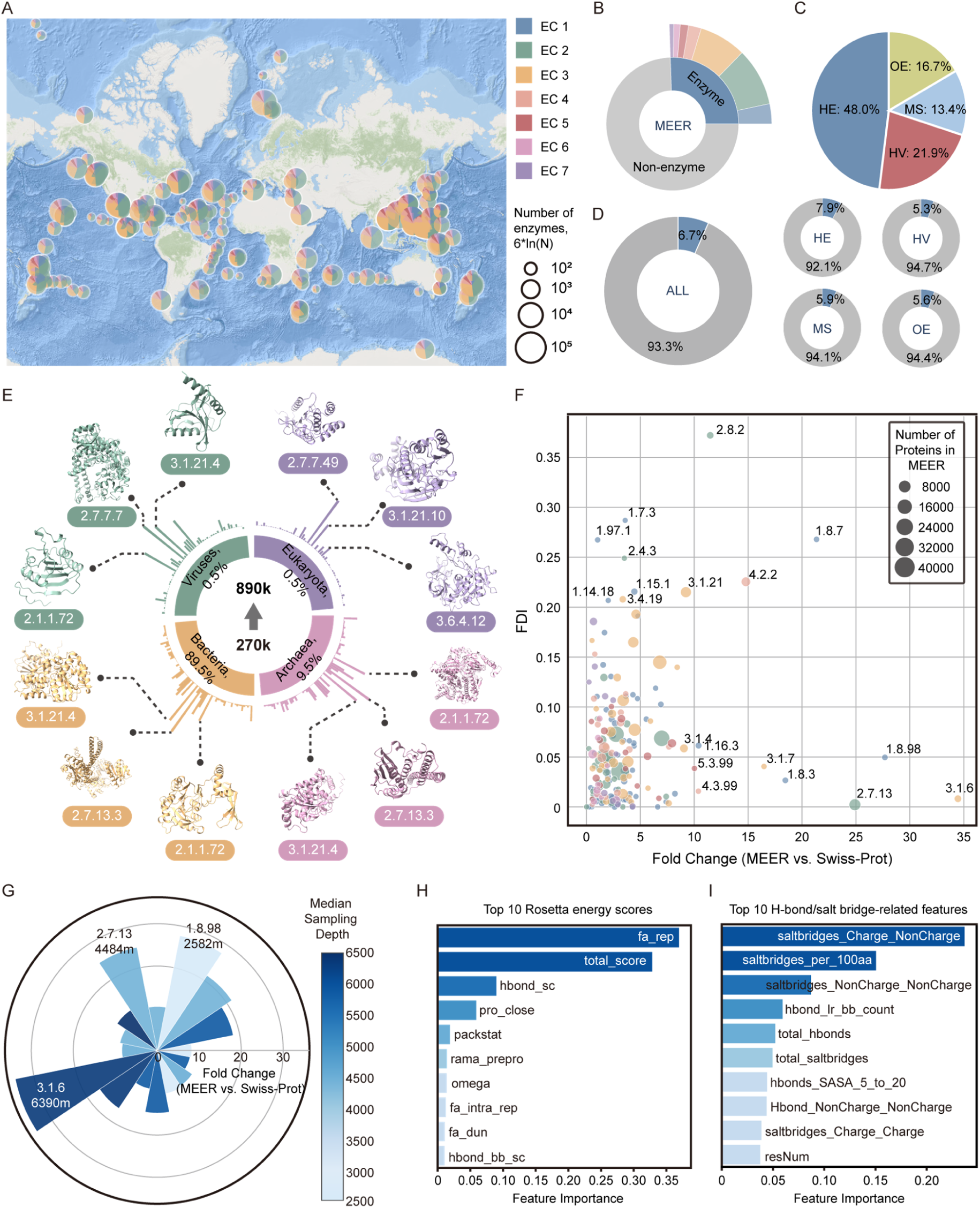
ACCESS unveils and characterizes an atlas of enzymes from extreme environments. (A) Global biogeography of novel enzyme classes. A world map illustrating the geographical distribution of predicted protein function enrichment across latitudes and longitudes, from the Arctic to the Antarctic, all major oceanic regions, and habitats ranging from shallow coastal areas to the Mariana Trench. Overlaid pie charts in each oceanic region show the proportional distribution of the seven major EC classes among newly discovered enzymes in each region, revealing large-scale biogeographical patterns in marine microbial metabolism. (B) Functional landscape of newly discovered deep-sea enzymes. The donut chart details the functional distribution of the 612,310 novel enzymes annotated by ACCESS from the MEER dataset. All seven major EC classes are represented, indicating a diverse metabolic potential in the deep sea. The most abundant classes are EC 2 (Transferases) and EC 3 (Hydrolases), highlighting their crucial role in a wide range of deep-sea metabolic and biogeochemical processes. The sheer volume and diversity of these findings represent a significant expansion of the known enzymatic universe. (C) Deep-sea habitats as hotspots for novel enzyme discovery. The chart quantifies the contribution of four distinct deep-sea habitat types to the discovery of novel enzymes. The abyssal zones (3,000–6,000 m depth) are the most prolific source, accounting for 48.0% of the new enzymes. Hydrothermal vents, cold seeps, and other habitats also contribute dramatically. This suggests that the unique and stable conditions of high pressure and low temperature in the abyss are potent drivers of novel protein function evolution. (D) Quantification of functional dark matter discovered by ACCESS. Donut charts depict the proportion of proteins newly assigned enzymatic functions by ACCESS, representing previously unknown functional dark matter. The overall proportion for the complete MEER dataset is shown, alongside the distribution across the four distinct habitats, revealing a substantial reservoir of novel enzymes (ranging from 5.30% to 7.90%) in each niche. (E) Taxonomic and functional landscape of newly discovered enzymes. The nested sunburst plot details the taxonomic origins and functional classes of the identified functional dark matter. The inner circle shows enrichment across four major taxonomic groups (Archaea: 9.50%, Bacteria: 89.5%, Fungi: 0.500%, Viruses: 0.500%). The outer circle displays representative 3D structures rendered in ChimeraX for the top three most abundant EC classes within each taxon, illustrating the structural diversity of the new atlas. (F) Comparative analysis of enzymatic functional space. A 2D scatter plot comparing the enzymatic functional space of the newly discovered MEER enzymes against the well-curated Swiss-Prot database. The x-axis represents the fold change in the number of enzymes for a given EC number (MEER vs. Swiss-Prot), showing functional expansion. The y-axis represents the Functional Diversity Index (FDI), showing sequence diversity within an EC class. Each point is an EC number, colored by its main class (1-7) and scaled by the total number of proteins. Points in the upper-right quadrant represent highly diverse and expanded functional families unique to extreme environments. (G) MEER-specific enzymatic functional sub-subclasses. A rose plot highlighting the 16 EC sub-subclasses (third-level EC numbers) with the highest fold enrichment, representing functions most specific to the MEER dataset. The length of each petal corresponds to the fold change value while the color intensity of each petal reflects the median sampling depth of enzymes in that subclass, confirming that these MEER-specific enzymes are not artifacts of shallow sampling but are robustly present in their native environments. (H) Structural stability features differentiating MEER enzymes. Bar charts showing the top 10 Rosetta energy score features identified by a random forest classifier as most predictive of enzyme habitat origin (MEER vs. Swiss-Prot). (I) Bar charts showing the top 10 H-bond/salt bridge-related features identified by a random forest classifier as most predictive of enzyme habitat origin (MEER vs. Swiss-Prot).

To quantitatively delineate the functional in the MEER-EC dataset, we plotted each EC sub-subclass by its enrichment in the MEER-EC dataset (fold-change relative to Swiss-Prot) against its content of functional dark matter indicated by the Functional Diversity Index (FDI) (Figures 3F, S3D). Analysis of the 16 sub-subclasses with the highest fold-change enrichment reveals a strong association with extreme depth, as their median sample origin depth far exceeds that of their Swiss-Prot homologs (Figure 3G). Among the most highly enriched functions were sulfuric ester hydrolysis enzymes like arylsulfatase (EC 3.1.6.1) and oxidoreductases acting on a sulfur group of donors such as sulfiredoxin (EC 1.8.98.2), suggesting that deep-sea ecosystems harbor unique biocatalytic solutions for recalcitrant substrate degradation and biological material synthesis under pressure ^65^. To quantitatively deconstruct the energy and biophysical features of deep-sea adaptation, we trained separate random forest models to classify proteins as originating from either MEER (extreme) or Swiss-Prot (non-extreme) datasets based on 30 distinct Rosetta energy score features (Figure 3H) and a set of 19 numeric features specifically quantifying the geometry and prevalence of intramolecular hydrogen bonds and salt bridges (Figure 3I). Both classifiers achieved high predictive accuracy in distinguishing deep-sea proteins from their terrestrial or surface-ocean counterparts. Subsequent feature importance analysis reveals the top energetic contributors. The most influential Rosetta-derived energy score feature was the Lennard-Jones atomic repulsive component (fa_rep). This term quantifies the energetic penalty of steric clashes and serves as an exquisite proxy for the quality and efficiency of hydrophobic core packing. This is consistent with the need to minimize pressure-induced water penetration into the protein core. It suggests that such optimized packing minimizes internal cavities, thereby resisting pressure-induced water penetration ^66–68^. Also ranking among the top three Rosetta-derived features was the side-chain hydrogen bond energy (hbond_sc), indicating the crucial contribution of extensive and optimized hydrogen bond networks in buttressing the protein fold against the immense extreme physical forces ^69^. Focused on electrostatic interactions in the second model, the overall density of intramolecular salt bridges emerged as a feature of importance. This finding corroborates our specific analysis of extremophilic polyethylene terephthalate hydrolases (PETases) in the next section and generalizes the principle that extensive, strategically distributed salt bridge networks are a cornerstone of protein stability under poly-extreme conditions ^70,71^.

Crucially, these dominant energy features, optimized hydrophobic packing, extensive salt bridge networks, and reinforced hydrogen bonding, are not merely statistical correlates, but constitute a coherent biophysical blueprint for conferring protein stability against hydrostatic pressure. Our analysis revealed convergent evolution at the energy level. When we examined functionally analogous enzymes (e.g., cellulases) sourced from phylogenetically distant microbial lineages within the MEER dataset, they consistently shared this same suite of distinguishing energy signatures even when their primary sequences showed little or no homology. This finding reinforces our hypothesis that energy features serve as conserved, function-linked evolutionary markers that transcend the limitations of conventional sequence and structural homology, thereby providing a powerful dimension for exploring the biosphere’s functional diversity.

### Experimental Validation Confirms ACCESS Predictions and Elucidates Adaptive Mechanisms

To ascertain the real-world predictive power of ACCESS, we established a high-throughput pipeline for the targeted synthesis of proteins and experimental validation of their functional annotations. The following case studies demonstrate how ACCESS not only correctly assigns function but also provides granular structural and energy insights necessary to explain the unique adaptive mechanisms of these novel enzymes.

Among the validated hits, ACCESS identified a protein, hereafter designated MEER-Chi1, as a chitinase (EC 3.2.1.14) with a prediction confidence of 85.7%. This protein represents a quintessential example of functional dark matter, which shares less than 0.250 sequence identity with any characterized chitinase in the Swiss-Prot database. Furthermore, its predicted structure exhibits a low TM-score of 0.566 when aligned to the closest annotated chitinase structure in the Swiss-Prot database (AFDB ID: AF-P11797-F1-model_v4). Despite this sequence and structural divergence (Figures S4A-S4H), the t-SNE plot generated from ACCESS’s multimodal embedding space shows that MEER-Chi1 clusters tightly with known, experimentally validated chitinases (Figures 4A, 4C). To understand the basis of its function, we performed a structural superposition of MEER-Chi1 with the aforementioned well-characterized bacterial chitinase ^72^ (Figure 4B). The analysis revealed a structurally conserved TIM-barrel catalytic core, including the canonical catalytic triad residues (D140, D142, and E144), which are essential for glycosidic bond hydrolysis ^73,74^. This conservation at the active site, despite widespread sequence divergence in surrounding regions (Figures 4D, 4E), explains how ACCESS correctly identified its function. Crucially, this superposition uncovered a distinct 71-residue C-terminal domain that is exclusively present in MEER-Chi1; no orthologous or paralogous counterparts of this domain were detected in other characterized homologs. This appended domain was delineated using the Encyclopedia of Domains (TED) computational pipeline ^75^, and subsequent annotation against the Pfam database classified it as a Secretion system C-terminal sorting domain ^76^. It implied MEER-Chi1 may exhibit dual functions, potentially coupling its canonical chitinolytic catalytic activity with a previously unrecognized secretion-associated role under the extremely high-pressure conditions of its deep-sea habitat ^77^.

**Figure 4.**
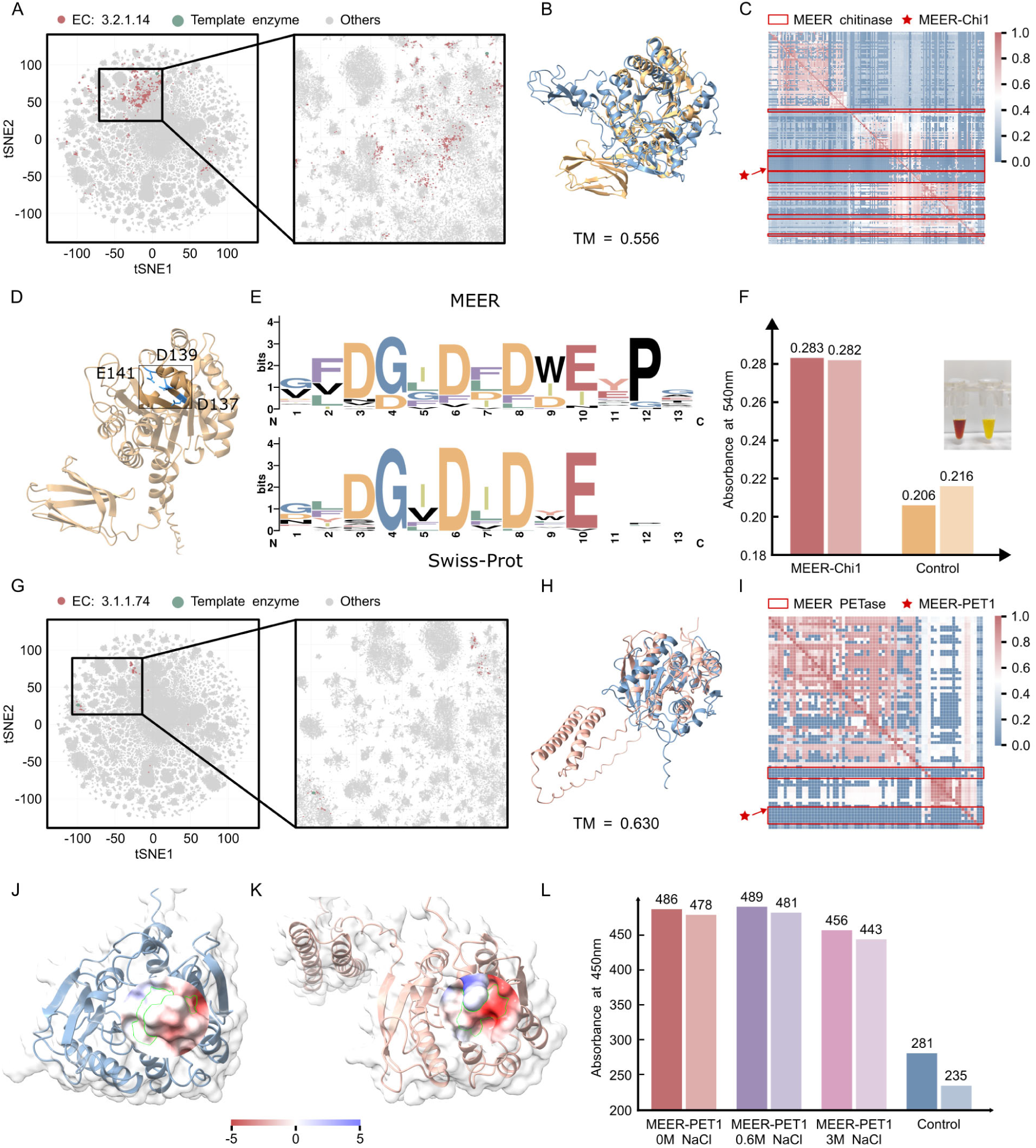
Experimental validation and mechanistic characterization of enzymes discovered from the MEER-EC database. (A) ACCESS’s multimodal embedding space separates novel and known chitinases. A t-SNE visualization of the embedding space for all proteins annotated with EC 3.2.1.14 (chitinase). This embedding is generated by ACCESS’s fusion of sequence, structure, and energy features. Each point represents a protein. Newly identified MEER chitinases (red dots, n=1,220) form a distinct, dense, isolated cluster from known chitinases in the Swiss-Prot database (green dot), predicting functionally distinctness of enzyme families. (B) Structural superposition reveals a novel domain. Structural superposition of a representative novel chitinase from MEER, MEER-Chi1 (yellow, marked with a red star in (C)) and a homologous template from the Swiss-Prot database (blue, UniProt ID: P11797) is shown. While the overall fold of the catalytic domain is conserved (RMSD = 2.6Å over 297Cα atoms), the MEER enzyme possesses a distinct novel C-terminal domain (approximately 71 amino acids). (C) Heatmap illustrates the pairwise sequence identity (lower triangle) and structural similarity (TM-score, upper triangle) among MEER-Chi1 and known chitinases (annotated as EC 3.2.1.14 in Swiss-Prot). The proteins are organized into three distinct blocks, revealing significant sequence and structural heterogeneity within this functionally related group. (D) Structure of a known chitinase. The known chitinase structure with its active sites (Asp140, Asp142, Glu144) is annotated. (E) Conservation of the catalytic triad. A sequence logo generated from a multiple sequence alignment of the catalytic domain MEER-Chi1 from panel c and the known chitinase from Swiss-Prot. Despite widespread sequence divergence across the domain, the analysis reveals strict conservation of the catalytic triad residues: Asp140, Asp142, and the catalytic nucleophile Glu144. (F) Experimental validation of chitinase activity. Enzymatic activity assay confirms the function of the recombinantly expressed and purified chitinase from MEER. The bar plot shows significant hydrolytic activity compared to an empty vector control, measured by the release of N-acetylglucosamine (absorbance at 540 nm). (G) A t-SNE plot of the ACCESS embedding space for PET-degrading enzymes. The MEER-derived candidates (red dots, n=111) form two distinct clusters. The investigation focuses on the cluster overlapping with the functional space of IsPETase (UniProt ID: A0A0K8P6T7) (green dot). (H) Structural superposition of a representative PET-degrading enzyme from the targeted cluster, MEER-Pet1 and IsPETase (green). The structures align well in the core catalytic region (RMSD = 3.62 Å over 208 Cα atoms); however, the MEER-Pet1 enzyme possesses a distinct novel N-terminal domain (approximately 53 amino acids). (I) A heatmap displays the pairwise sequence identity and structure similarity (TM-score) for the newly predicted MEER PET-degrading enzymes. Similar to chitinases, these enzymes exhibit low sequence and structural conservation, organizing into two primary blocks that correspond to the clusters observed in panel f. This again highlights the diversity of functional solutions evolved for the same catalytic task. (J-K) Comparison of the 3D electrostatic potential surface between IsPETase (J) the MEER-Pet1 (K) reveals a more pronounced negative electrostatic potential within the substrate-binding cleft of MEER-Pet1. (L) Enzymatic activity assay of the recombinantly expressed novel PETase, tested under varying salinity conditions (0 M, 0.6 M, and 3.0 M NaCl). The enzyme demonstrates robust activity against a bis(2-hydroxyethyl) terephthalate (BHET) substrate (absorbance at 450 nm). While its activity decreases with increasing salt concentration, the enzyme retains substantial catalytic function even at 3.0 M NaCl, demonstrating strong halotolerance.

Despite widespread sequence divergence across the domain, the analysis of the sequence logo generated from a multiple sequence alignment of the catalytic domain reveals the strict conservation of the catalytic triad residues (Figure 4E). To experimentally validate this prediction and investigate its predicted piezophilic nature, we conducted a series of in vitro enzymatic assays to measure the activity of purified MEER-Chi1 against a colloidal chitin substrate. The results from two independent biological replicates, when compared against an empty-vector control group, revealed a statistically significant increase in hydrolytic activity, confirming the protein’s predicted chitinase function (Student’s t-test, p < 0.01; Figure 4F).

In another case of functional discovery, ACCESS assigned PETase (EC 3.1.1.74) to a previously uncharacterized protein named MEER-Pet1 with a confidence score of 0.829. Similar to the chitinase, MEER-Pet1 is sequence dark matter, with < 0.250 identity with the canonical PETase from *Ideonella sakaiensis* (PDB: 6EQE) ^78^ (Figures 4G, 4I). This substantial sequence divergence underscores the limitations of homology-based functional inference (Figures S4I-S4P). The t-SNE visualization showed that the MEER-derived candidates (red dots) formed two distinct clusters, indicating potential functional or structural sub-specialization (Figure 4G). Despite the low global similarity, the MEER-PET1 model retains a canonical Ser-His-Asp catalytic triad (S226, H279, D307) (Figure 4H), which is the essential chemical machinery for ester bond hydrolysis ^79,80^. Notably, MEER-Pet1 exhibits a previously unreported N-terminal domain of 53 amino acids. Although this TED-predicted domain currently lacks functional annotation in the Pfam database, it offers critical insights that could guide rational protein design efforts aimed at optimizing PET hydrolases ^81,82^.

Beyond static tertiary structure, the functionally salient attribute of the MEER enzymes is their uniquely tailored surface bioelectric landscape. Our analysis reveals a more negative electrostatic potential within the substrate-binding cleft of MEER-Pet1. We postulate that this charged environment functions as an electrostatic funnel, actively steering the electrophilic carbonyl carbons of the PET polymer’s ester linkages into the active site. This electrostatically guided mechanism would optimize substrate orientation and accelerate catalytic turnover ^83,84^ (Figures 4J, 4K). While a negatively charged surface might seem counterproductive for binding PET, which can develop negative charges during hydrolysis, we hypothesized that this electrostatic architecture is not a primary catalytic adaptation but rather a fundamental evolutionary solution for extreme halotolerance ^85,86^. In high-salinity environments, a dense negative surface charge helps maintain a stable hydration shell by coordinating cations, preventing protein aggregation and denaturation. This solvation shield likely maintains the enzyme’s structural integrity and solubility, which is a prerequisite for any catalytic activity in a high-salt milieu. To test this hypothesis, we synthesized the gene for MEER-Pet1, expressed the protein heterologously in *E. coli*, and assayed its PET degradation activity across a range of NaCl concentrations (0.1 M, 0.6 M, and 3.0 M), using amorphous PET film as the substrate. The results, analyzed by a two-way Analysis of Variance (ANOVA), confirmed our in-silico hypothesis (Figure 4L). In contrast to the empty vector control, MEER-Pet1 retained its degradation activity at 3.0 M NaCl, demonstrating it is not merely halotolerant but is a true haloduric enzyme.

### ACCESS’s Interpretability Pinpoints Functionally Critical Active and Allosteric Residues

A central challenge in harnessing the functional potential of the uncharacterized protein dark matter from unique environments is to move beyond mere functional annotation to a mechanistic understanding that enables rational engineering. To address this, we probed ACCESS’s residue-level interpretability, converting its black-box EC predictions into an *in silico* mutagenesis platform for rational design (Figure 5A). We quantified each residue’s contribution to the final EC number prediction of a protein by fusing gradients from the node and its neighbors during ACCESS’s back propagation ^87^. In essence, residues with high scores are those that the model identified as most influential in determining the protein’s catalytic function. To validate the biological relevance of these importance scores, we benchmarked the module’s performance on well-characterized enzymes from the Swiss-Prot database with experimentally annotated active and binding sites. For each enzyme, we ranked its residues by their ACCESS-generated importance scores and assessed whether the top-ranked residues corresponded to known functional sites. The results demonstrated remarkable predictive power: ACCESS correctly identified active site residues with over 0.800 accuracy for 64.3% of the enzymes tested (Figure 5B) and, even for the more conformationally flexible and diverse binding sites, maintained over 0.800 accuracy for 22.5% of the enzymes.

**Figure 5.**
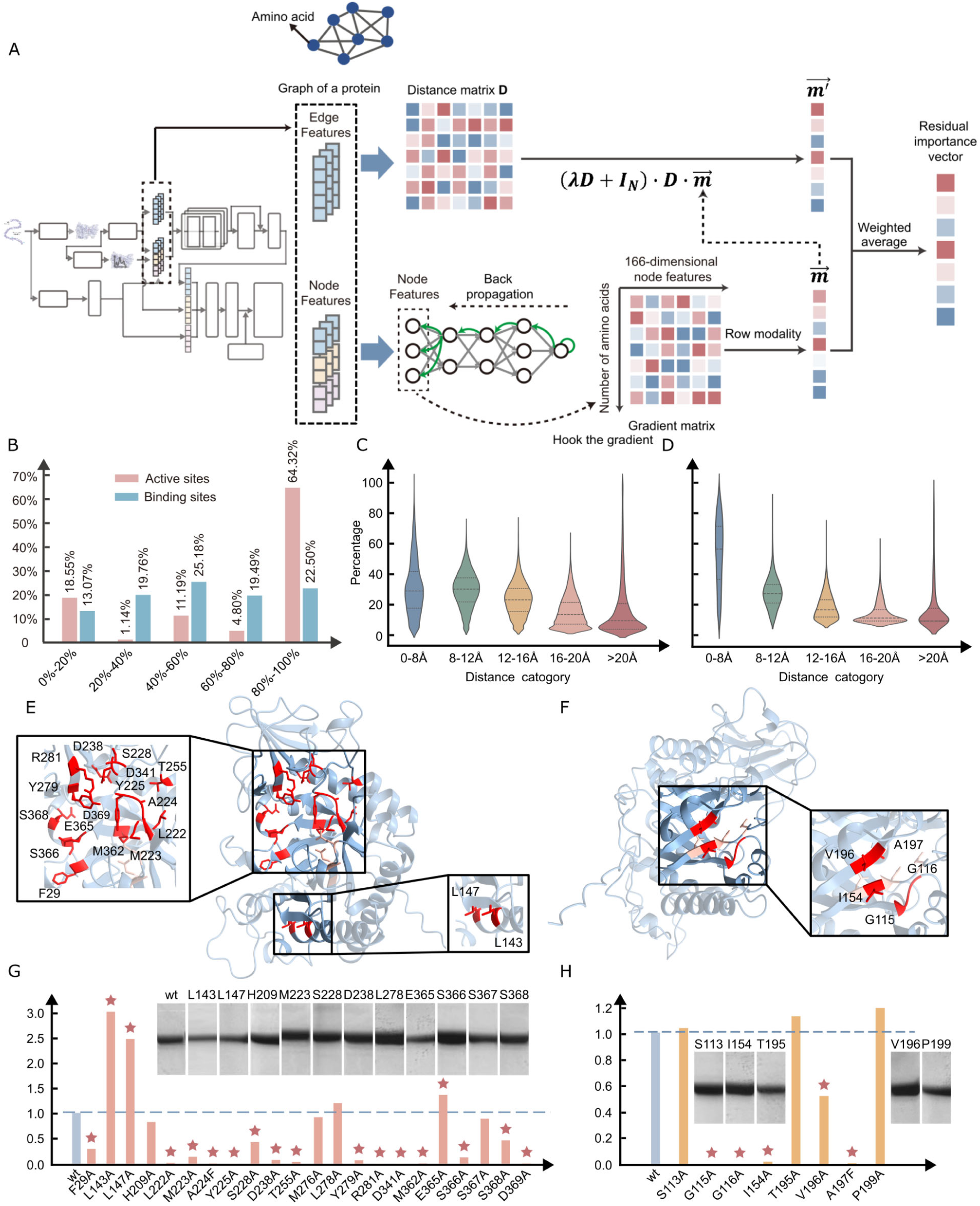
Interpretability of ACCESS Enables Residue-Centric Rational Design of Protein Function. (A) The flowchart depicts the architecture of our post-hoc traceability mechanism, an integrated gradient-based saliency mapping framework specifically adapted for our multimodal MPNN. The process begins with a protein’s multimodal graph representation, which encodes sequence, structure, and energy information. After ACCESS predicts the function, the mapping algorithm computes the gradient of the cosine similarity between the predicted and real label embedding. This process generates a saliency score for every residue, quantifying its contribution to the predicted function. Residues with the highest saliency scores are identified as functionally critical. A key technical innovation demonstrated here is the development and application of a sophisticated post-hoc traceability mechanism to our multimodal MPNN architecture, enabling the deconstruction of a complex “black box” prediction into an interpretable, residue-level functional attribution map. (B) We benchmarked ACCESS’s saliency predictions against all enzymes with experimentally annotated active sites and binding sites. The x-axis categorizes the sites into ‘Active Sites’ and ‘Binding Sites’, while the y-axis represents the percentage of enzymes for which our method achieves high prediction accuracy. We define a stringent accuracy metric where a prediction is considered successful if over 80.0% of the annotated site residues are captured within the top-ranked salient residues predicted by ACCESS. The results show that for 64.3% of all annotated enzymes, ACCESS successfully identifies the active site with this high degree of accuracy. Similarly, high-accuracy prediction of binding sites is achieved for 22.5% of the relevant proteins. (C-D) These violin plots offer a nuanced analysis of the spatial distribution of the critical residues predicted by ACCESS, relative to known functional centers. For the same Swiss-Prot enzyme dataset, we calculated the minimum Euclidean distance from each high-saliency residue to the nearest atom of an annotated active site (C) and binding site (D). The width of the violins represents the density of predicted residues at a given distance. A significant population of predicted residue clusters at very short distances (< 5 Å), corresponding to ‘on-site’ residues directly within or adjacent to the functional pocket. Crucially, the plots also show another substantial population of predicted residues located at significantly long distances (10–30 Å or more), representing putative allosteric sites. (E-F) These two figures present 3D structural representations of the known chitinase PoChi, providing a spatial context for the experimentally validated results from panels (G) and (H). The protein backbone is shown as a cartoon, with the catalytic active site highlighted in pink. In both structures, the 23 functionally critical residues validated by alanine scanning are rendered as sticks. The left structure (E) highlights the 18 validated allosteric residues, illustrating their distributed locations on the protein surface, distant from the active site. The right structure (F) exclusively highlights the 5 validated ‘on-site’ residues, showing their close proximity to the active site, consistent with their direct roles in substrate binding or catalysis. This panel’s key contribution is the 3D visualization of functionally validated control points, providing an actionable blueprint for targeted protein engineering. This visual evidence bridges the gap between abstract saliency scores and concrete structural locations, empowering researchers to rationally design proteins with modified activity, stability, or specificity by targeting either local or long-range functional hotspots. (G-H) Direct experimental validation and the expressed SDS-PAGE image of both on-site residues (G) and allosteric residues (H) for the predictions made by ACCESS using chitinase from PoChi as a model system. Based on the saliency map generated for chitinase, we selected the top 30 predicted critical residues for experimental characterization via site-directed alanine scanning mutagenesis. The bar charts display the relative catalytic activity of each mutant enzyme compared to the wild-type samples, measured using a fluorogenic substrate assay. The results strikingly confirm our predictions: 23 out of 30 mutations led to a significant impact (stars, green stars denote weak or non-expression) in enzyme activity. Of these, 5 were on-site residues located within the substrate-binding cleft, while the other 18 were distal allosteric residues, some over 20 Å away from the active site, thereby validating ACCESS’s capability to identify both direct and long-range functional residues.

The interpretability of the ACCESS framework lies beyond the conventional focus on the immediate catalytic environment. It also identified residues of high functional importance located at significant distances from the active site, flagging them as potential allosteric control points (Figures 5C, 5D). We defined these as "allosteric hubs": amino acids that, despite being spatially distal (> 8 Å from any catalytic residue), receive high attention weights from the model. Such sites are notoriously difficult to predict ^88,89^. ACCESS enables the deconvolution of predicted critical residues into two distinct populations: direct on-site catalytic residues and distal allosteric hubs. This capability represents a paradigm shift from conventional methods based on sequence or structure alignment, which are inherently biased toward highly conserved residues within the catalytic pockets and often fail to capture the subtle, long-range interactions that govern allosteric regulation.

In a case study of the well-characterized chitinase, PoChi (GenBank: PQ195456) ^90^, ACCESS computationally scored the 30 most functionally significant residues in PoChi, comprising 8 on-site catalytic residues and 22 putative allosteric hubs. Among the latter, Aspartate-341 (D341) emerged as a top-ranked hub despite lying 23Å away from the catalytic Glutamate 157 (E157) (Figure 5E). Guided by these predictions, we computationally identified the 30 most functionally significant residues in PoChi, comprising 8 on-site catalytic residues and 22 putative allosteric hubs, including D341. We then experimentally validated these predictions using alanine scanning mutagenesis, a systematic method where target residues are replaced by alanine to eliminate side-chain interactions and thereby probe their functional contribution. We generated single-point mutants for the top predicted residues and measured their enzymatic activity relative to the wild-type (Figures 5G, 5H). As expected, mutating the top-ranked active site residues, such as A197F and G115A, resulted in a near-complete loss of function, with measured activity dropping by over 98.0%. This validated the model’s ability to identify the core catalytic machinery. More revealingly, mutations at predicted allosteric sites produced diverse and significant effects. The D341A mutant exhibited a 65.0% reduction in activity, confirming its crucial role in maintaining the structural integrity required for catalysis, likely by disrupting domain stability under assay conditions ^74^. To be noted, two of the allosteric mutations (L143A, L147A) even cause more than a 2.5-fold increase in activity.

Mapping the 10 ACCESS-prioritized residues on the 3D structure of MEER-Chi1 provided a clear structural rationale for their predicted functional contributions (Figures 5E, 5F). The five catalytic residues at the active site formed a dense, geometrically optimized constellation within the enzyme’s active site cleft, poised for substrate binding and catalysis. In contrast, the five predicted allosteric regulators, including D341 and A224, were dispersed across the protein surface, forming a distal network. This clear spatial segregation, corroborated by the experimental mutation data, validates ACCESS’s power to generate actionable hypotheses for protein engineering. By focusing experimental efforts on a small, high-potential subset of residues, this predict-design-test workflow can reduce the screening burden associated with random mutagenesis or full saturation scanning by over 90.0%, dramatically accelerating the discovery and optimization of novel enzymes from functional dark matter.

### ACCESS Reveals a Functional Phylogeny Across the Tree of Life

To investigate the macroevolutionary patterns of protein function beyond the constraints of sequence homology, we applied ACCESS to the predicted proteomes of 48 phylogenetically diverse species from the AlphaFold Database (AFDB) ^49^. This comprehensive analysis, spanning archaea, bacteria, fungi, protists, plants, and animals, enabled the construction of a vast functional encyclopedia based on enzymatic capabilities (EC numbers). Our results reveal a landscape of function-driven, energy-guided evolution, where an organism’s enzymatic repertoire is a direct reflection of its ecological niche and metabolic strategy. This functional narrative often diverges significantly from those based on traditional, sequence-based phylogenies ^91^, underscoring the unique power of our energy-centric approach.

We first constructed a comprehensive, phylogenetically resolved enzymatic cartography spanning the tree of life, with ACCESS assigning specific EC numbers to 673,560 proteins across the 48 representative species. Projecting this vast functional dataset onto a phylogenetic scaffold provided a panoramic view of enzymatic distribution across diverse evolutionary lineages (Figure 6A). This cartography highlighted species-specific functional biases, reflecting distinct evolutionary trajectories and adaptive pressures. For instance, our analysis captured the dramatic expansion of the protein kinase repertoire (EC 2.7.11.1) associated with the prokaryote-eukaryote transition, a watershed event in the history of life. While the proteome of the bacterium *E. coli* contains a modest 12 distinct kinase variants, the unicellular eukaryote *Saccharomyces cerevisiae* possesses 47. This nearly fourfold diversification mirrors the evolutionary leap in cellular complexity of eukaryotes. These organisms utilize extensive phosphorylation-based signaling cascades to regulate virtually all cellular processes, from cell cycle progression to environmental stress response ^92–95^. Furthermore, our analysis uncovered a unique functional signature within the animal kingdom, revealing a high diversity of trypsin-like serine proteases (EC 3.4.21.4) in *Drosophila melanogaster*. ACCESS identified 32 distinct functional variants of this enzyme class, greater than in any other analyzed animal species, including vertebrates. This expansion of the protease family correlates with its digestive physiology, adapted to a diet of fermenting fruit rich in microbial and plant proteins ^96–100^. To substantiate the predictive fidelity of this enzymatic map, we conducted a rigorous, multi-faceted validation of ACCESS’s performance against the manually curated Swiss-Prot database. The high scores across all six key performance metrics for the complex proteomes of *H. sapiens, D. melanogaster* and the other three species underscore ACCESS’s ability to assign not only correct functions but also their precise positions within the intricate enzyme classification system (Figures 6B, S5A). The consistently high accuracy and F1-scores across both non-enzymatic proteins and all seven main EC classes confirm the unbiased nature of ACCESS’s high prediction accuracy (Figures 6C, S5B). This is further illustrated by an investigation of 11 human proteins from the same enzyme class, alpha-1,4-glucan branching enzyme (EC 2.4.1.41). The plot of sequence and structural space reveals significant divergence in both sequence identity and TM-score (Figures 6D, S5C). Among them, the 3D structural models of eight representative proteins show structural variations ranging from compact globular domains to more complex, multi-domain architectures (Figures 6E, S5D).

**Figure 6.**
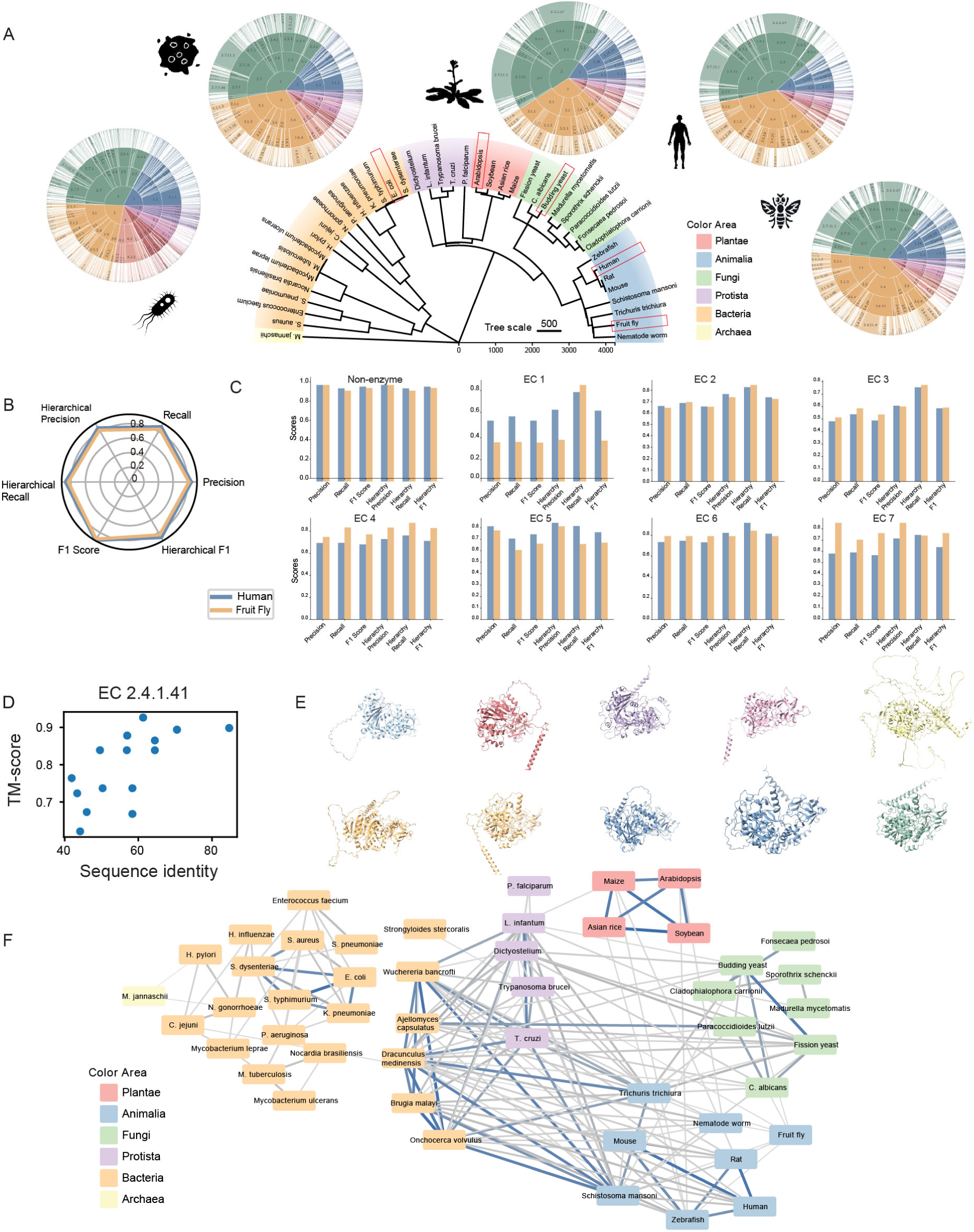
ACCESS Reveals Macroevolutionary Trends and Functional Phylogenomics Across the Tree of Life. (A) A semi-circular phylogenetic tree delineating the evolution of enzymatic functional diversity across five key species representing major evolutionary nodes: *Escherichia coli* (prokaryote), *Saccharomyces cerevisiae* (fungal eukaryote), *Drosophila melanogaster* (animal eukaryote), *Arabidopsis thaliana* (plant eukaryote), *and Homo sapiens* (mammalian eukaryote). Adjacent circular heatmaps depict the functional diversity profile, defined as the number of distinct proteins assigned to each terminal EC number. Comparative analysis shows an expansion in the diversity of non-specific serine/threonine protein kinases (EC 2.7.11.1) and ubiquitin-conjugating enzymes (EC 2.3.2.27) from prokaryotes to eukaryotes. *D. melanogaster* exhibits an unparalleled diversification of trypsins (EC 3.4.21.4). (B) A radar chart quantifying the consistency between functional predictions generated by ACCESS and annotations from the gold-standard resource, the Swiss-Prot database. The chart evaluates six key performance metrics for proteins from *H. sapiens* and *D. melanogaster*, including precision, recall, F1-score, hierarchical precision accounting for EC hierarchy, hierarchical recall, and hierarchical F1-score. (C) Performance metrics for non-enzymatic and all seven main EC classes of proteins from *H. sapiens* and *D. melanogaster*, confirming the unbiased nature of ACCESS’s high prediction accuracy. (D) Distribution of 11 human proteins assigned to EC 2.4.1.41 (alpha-1,4-glucan branching enzyme) exhibiting divergence in sequence identity and TM-score (a measure of structural similarity). (E) Visualization of 3D structural models of eight representative proteins from the EC 2.4.1.41 set in (D) with structural variation. (F) Visualization of the functional relationships among all 48 model species using a force-directed network graph. Each node in the network is a species while the edges represent enzymatic functional profiles similarity greater than 0.900, with thickness proportional to similarity. Blue edges highlight similarities > 0.970. The network shows remarkable congruence with the canonical tree of life, with species from Archaea, Bacteria, Fungi, Protista, Plants, and Animals forming distinct clusters. Bacteria are bifurcated into two superclusters: one is functionally more like Archaea; the other is closer to protists and animals. The parasite *Schistosoma mansoni* clusters with bacteria. Plants form a highly cohesive and isolated cluster. Cosine similarity scores > 0.950 indicate high functional similarity among humans, mice, and rats.

Zooming out from individual enzyme classes, we compared the overall functional similarity between organisms. We vectorized each species’ complete EC profile and computed a pairwise cosine similarity matrix, which was then visualized as a force-directed network (Figure 6F). In this network, nodes represent species, and edges represent their functional similarity, offering a global view of evolutionary relationships through a functional lens. The resulting network self-assembles into kingdom-spanning continents where archaea, plants, fungi and animals collapse into tight, blue-edged clusters (internal similarity > 97.0%), mirroring textbook phylogeny and confirming that ACCESS’s EC profiles are a quantitative proxy for evolutionary kinship.

The network’s real power, however, is to reveal functional convergences and divergences that transcend traditional genomic or genetic sequence-based taxonomy. For instance, the bacterial domain, which appears monophyletic in sequence-based phylogenetic trees, is split into two distinct functional clades. One showed high functional similarity to archaea, while the other clustered more closely with protists. This split suggests that metabolic strategy and environmental adaptation are powerful evolutionary forces capable of forging functional relationships transcending phylogenetic boundaries ^101^. This principle of function over phylogeny was further exemplified by the parasitic flatworm *Schistosoma mansoni*. Despite being an animal, its functional profile showed 0.650 similarity to bacteria, far exceeding its 0.400 similarity to other metazoans in our dataset. This seemingly paradoxical clustering reflects the parasite’s profound metabolic dependence on endosymbiotic bacteria ^102^. Finally, the network provided a quantitative validation for the use of model organisms in biomedical research. *Homo sapiens*, *Mus musculus*, and *Rattus norvegicus* formed a tight, distinct cluster with over 0.950 functional similarity. This high degree of conservation in their core enzymatic toolkits provides a quantitative, function-level justification for the utility of rodent models in studying human metabolism and disease ^103,104^.

## DISCUSSION

This work introduces a paradigm shift in functional genomics by leveraging the conserved energetic blueprints to decode the vast dark matter of extremophile proteins. Conventional computational tools face a fundamental limitation: divergent evolution erases the sequence-level signals of common ancestry upon which they rely. While recent AI methods have begun to overcome this by learning complex sequence patterns ^18^, they largely ignore the physical principles governing protein function. ACCESS bridges this critical gap through its multimodal integration of energy landscapes, evolutionarily constrained features that directly shape protein action and stability. Our results demonstrate that ACCESS establishes a more robust and generalizable foundation for function prediction, outperforming SOTA tools by over 20.0% on non-homologous protein sets.

The construction of the MEER-EC functional atlas showcases the discovery power of our energy-centric, multimodal AI approach. By computationally characterizing nearly one million previously unknown enzymes from deep-sea environments, we have generated a resource that dramatically expands the known functional enzyme universe and provides a quantitative framework for understanding molecular-level adaptation ^44^. Notably, 92.2% of the cataloged enzymes are newly discovered, with 33.7% sourced from hadal zones such as the Mariana Trench. This repository includes numerous high-value candidates for industrial and environmental applications, such as haloduric PETases for degrading plastics in high-salinity environments and pressure-resilient chitinases for biowaste conversion ^105^. This efficient *in silico* bioprospecting addresses key limitations of traditional marine discovery, which is often hampered by high capital costs, low success rates, and unsustainable harvesting practices ^106^. Our findings position energy minimization as a universal selective driver in extremophile evolution. Across vast phylogenetic distances, proteins from the repository exhibit conserved energetic features, such as decreased salt-bridge networks and optimized charge distributions. This phenomenon explains the observed energetic convergence, whereby the recurrence of optimal energetic signatures in functionally analogous enzymes from phylogenetically distinct organisms challenges the longstanding paradigm that sequence homology is the primary indicator of functional relatedness. Beyond its predictive prowess, ACCESS is not a "black box". It offers critical interpretability, transforming it from a prediction oracle into a causal inference engine for protein engineering. By mapping specific energetic features to catalytic and allosteric sites, ACCESS provides both mechanistic insights and rational design targets of a well-characterized chitinase. This capability moves the field beyond laborious trial-and-error screening towards a hypothesis-driven, AI-guided design cycle.

ACCESS’s functional encyclopedia further uncovers novel evolutionary mechanisms in key organisms. For instance, the bacterial-like EC profile in *Schistosoma mansoni* suggests a history of endosymbiosis-driven adaptation, where hardy enzymatic functions were likely acquired through horizontal gene transfer. Similarly, the unique protease diversity in *Drosophila* species reflects niche-specific adaptations encoded in energetic stability. These insights prompt a redefinition of protein evolution as a function-first process: organisms adapt to extreme environments primarily through functional shifts that are directly encoded in energetic landscapes, rather than through the accumulation of sequence mutations alone ^107–109^. Furthermore, extending this framework to entire proteomes of key organisms provides a new functional-evolutionary lens to reinterpret the tree of life, allowing us to track the birth, death, and transformation of molecular functions. The integration of these multiple data modalities paves the path toward a more holistic understanding of biological systems ^110^. Looking forward, ACCESS establishes a foundational framework for the next generation of functional genomics and intelligent bioprospecting. As both a predictive tool and an analytical platform, ACCESS is poised to systematically accelerate the discovery of functional protein dark matter, enabling a more sustainable and efficient exploration of the biosphere’s most enigmatic frontiers.

### Limitations of the study

While ACCESS represents a leap forward, its reliance on computationally derived energy landscapes presents a primary challenge at the intersection of high-performance computing and biophysical AI. The substantial computational cost of accurately calculating structure and energy functions, particularly for large proteins or complex folds, remains a bottleneck for widespread accessibility. Additionally, achieving highly accurate protein structures remains a critical objective although we have implemented optimized molecular dynamics relaxation protocols and Rosetta FastRelax ^111^ to optimize the predicted structures. A second constraint relates to sampling biases inherent in the MEER-EC Atlas. The formidable challenges of deep-sea exploration have resulted in a geographically fragmented data landscape, where accessible hadal zones are disproportionately overrepresented while vast abyssal and ultra-abyssal plains remain critically undersampled. Expanding partnerships with international oceanographic initiatives is therefore crucial for achieving comprehensive coverage. Further consideration involves experimental validation, which was conducted primarily on recombinant proteins expressed in model systems. These conditions may not fully replicate the native physicochemical milieu of extremophiles, potentially altering enzyme kinetics or stability. Future studies should prioritize *in vitro* assays under simulated deep-sea conditions, such as high hydrostatic pressure and low temperature, to confirm functional predictions in ecologically relevant contexts. Finally, while ACCESS outperforms SOTA tools on non-homologous datasets, its performance on certain enzyme classes, particularly those reliant on complex cofactors, remains suboptimal. This likely suggests that energy-based features may not capture all aspects of enzyme mechanism, such as cofactor-dependent catalysis. Incorporating additional modalities, such as quantum mechanical descriptors of reaction chemistry, represents a promising direction to address these gaps and further enhances the model’s predictive power.

Despite these limitations, ACCESS establishes a robust foundation for decoding the functional repertoire of extremophile proteins. Its multi-modal deep learning framework provides a scalable and adaptable approach for exploring functional dark matter, with important implications for biotechnology, evolutionary biology, and ecological conservation.

## ACKNOWLEDGMENTS

We thank Prof. Xiang Xiao from Shanghai Jiao Tong University for the helpful discussion and critical reading of the manuscript. This work has been supported by the National Key Research and Development Program of China (2025YFC2817100), the National Natural Science Foundation of China (32225039, 32100514, and U21A20271), the Guangdong Natural Science Funds for Distinguished Young Scholar (2025B1515020038), the Taishan Scholar Project of Shandong Province (tstp20250748, tsqn202312099), the Shenzhen Science and Technology Program (JCYJ20250604191305008), the Major Science and Technology Research Project of Sanya Yazhou Bay Science and Technology City (SKJC-2021-01-002), the Major Scientific and Technological Innovation Project of Shandong Province (2024ZLGX07), the Central Government Guides Local Science and Technology Development Fund for Xinjiang (ZYYD2025ZY01), the Agriculture Research System of China (CARS-48), the Natural Science Foundation of Shandong Province (ZR2024Y0054), and the Support Program for Youth Innovation Technology in Colleges and Universities of Shandong Province (2023KJ041). We acknowledge the support from the China National GeneBank data storage and database construction.

## AUTHOR CONTRIBUTIONS

X.M., G.F., and S.M.Y.L. conceived and directed the project. M.X., J.W., H.Y., X.X., Y.S., and X.F. designed experiments and analyses. M.X., D.T.W., H.J., and Q.L. developed the ACCESS method. Q.L., D.T.W., X.L., and Y.L. collected and analyzed the data. H.J., D.W., H.D., H.P., Y.N.Z., J.C., and X.W. performed functional experiments. X.Y., Y.J.L., A.X., and Y.W. performed database construction. S.Y.L., H.L., D.L., S.S.L., L.M., Y.X.L., C.X., L.J., Y.M.Z., J.S., M.W., Y.G., Z.L., T.M., Y.Z., and C.H.X. assisted in the data analysis. M.X. drafted the manuscript with input from all authors. D.T.W., H.J., X.L., Y.L., G.F., and X.M. edited the manuscript.

