## Supplemental Materials for "Multimodal AI Decodes Extreme Environment Functional Dark Matter Beyond Homology"

### SUPPLEMENTAL INFORMATION

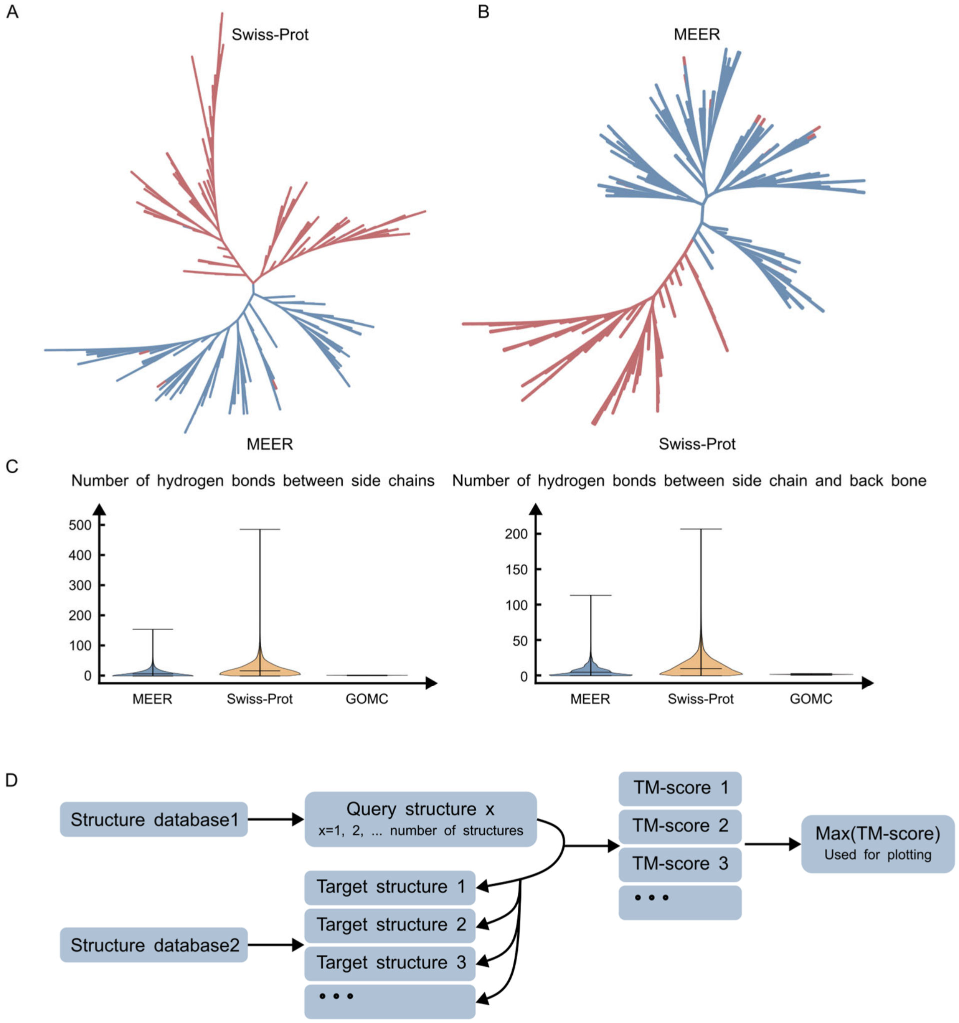


**Figure S1. Additional comparison between MEER, Swiss-Prot, and GOMC databases, related to Figure 1.**

(A-B) Maximum-likelihood phylogenetic trees constructed using sequences with similar structures compared to structures with AFDB ID: AF-F0XS04-F1-model_v4 and AF-B5FYY5-F1-model_v4, respectively. The clustering results also show distinct divergence with only a few exceptions.

(C) Violin plots showing the number of hydrogen bonds between side chains (left) and the number of hydrogen bonds between side chains and backbone (right).

(D) Pipeline of computing structure similarity between databases. Query structures from database 1 are compared to all the target structures from database 2 sequentially. And then the maximum of the computed TM-scores is used for plotting.

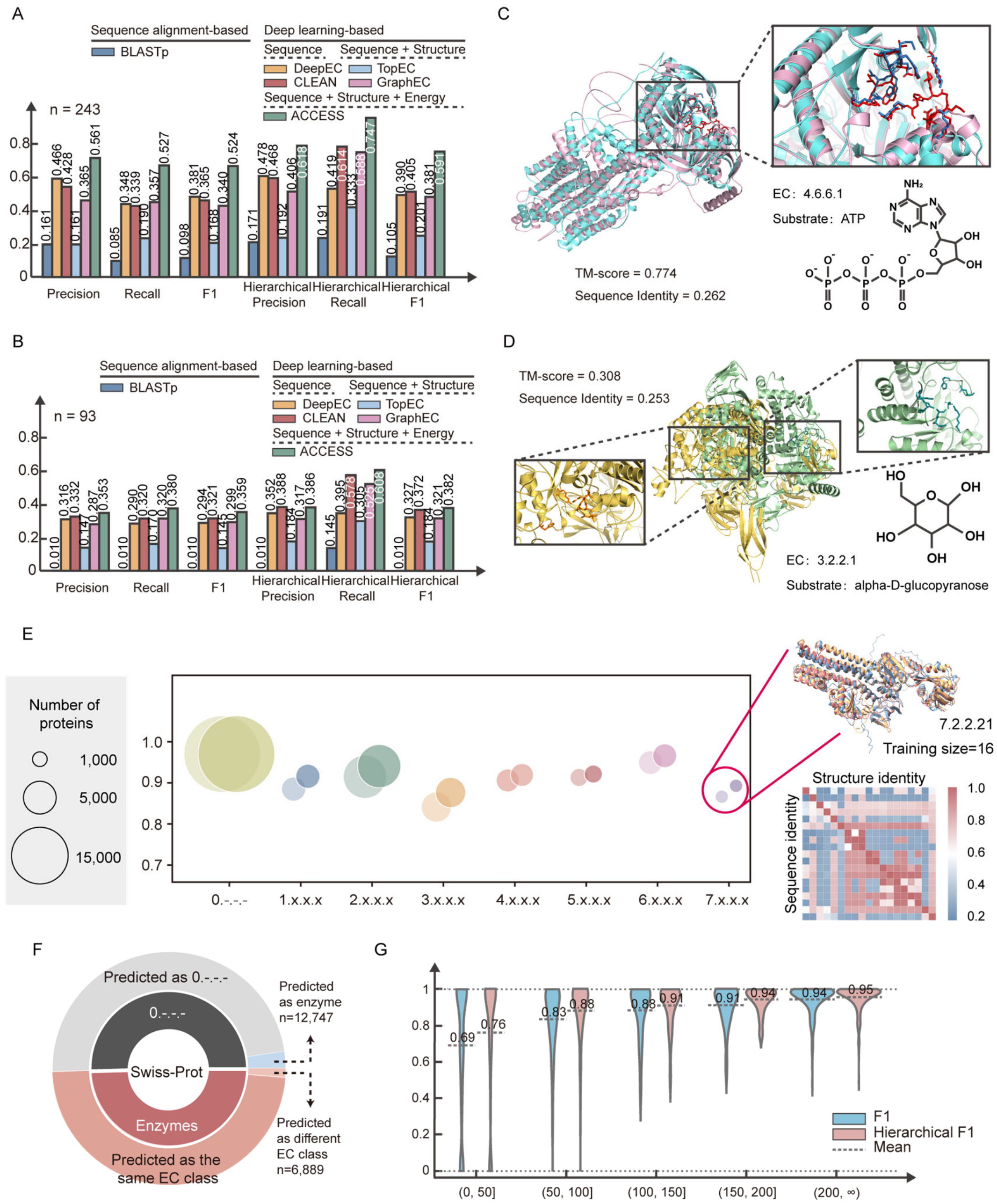


**Figure S2. Additional benchmark performance and application of ACCESS, related to Figure 2.**

(A) ACCESS demonstrates robust performance on proteins with low sequence identity. The bar chart benchmarks existing tools on a specially curated dataset which contains 243 proteins with less than 0.300 sequence identity to any protein in the training data. This test is designed to assess a model's ability to predict function beyond sequence homology. ACCESS maintains its superior performance in terms of all six metrics.

(B) ACCESS demonstrates robust performance on low-homology "functional dark matter" proteins. This bar chart benchmarks the same set of tools on the specially curated spDarkM-93 dataset, which contains proteins with less than 0.300 sequence identity and less than 0.500 structure similarity to any protein in the training data. This test is designed to assess a model's ability to predict function beyond homology. ACCESS maintains its superior performance in terms of all six metrics, while the performance of other tools, which often rely implicitly or explicitly on sequence or fold similarity, degrades significantly.

(C) Case study of function prediction beyond sequence homology. ACCESS identifies two enzymes as Type VI adenylyl cyclases (EC 4.6.1.6) despite their low sequence identity, demonstrating its power to detect shared function in the absence of clear sequence homology.

(D)Case study of function prediction beyond homology. This panel illustrates the key strength of ACCESS with a real-world example: two enzymes that share the identical function of purine nucleosidase (EC 3.2.2.1). Despite their identical catalytic activity, these proteins exhibit distinct sequence and structure divergence.

(E) Performance of ACCESS across non-enzymes and the seven EC classes. Bubble pairs show F1-scores (light tone) and hierarchical F1-scores (dark tone) for each class; bubble area is proportional to the number of enzymes in that class. The right-hand panel illustrates EC class 7 with a concrete example: EC 7.2.2.21 is represented by only 16 training samples, yet ACCESS attains an F1-score of 1.0. The bottom-right heatmap displays the pairwise sequence and structural similarity among these 16 proteins.

(F) Large-scale curation and refinement of public protein databases using ACCESS. ACCESS provided new enzymatic function annotations for 12,747 proteins that were previously annotated merely as "hypothetical" or were presumed to be non-enzymatic, thereby illuminating new functional components of the proteome. Furthermore, it corrected 6,889 existing annotations that were inconsistent with its high-confidence predictions.

(G) Distribution of F1-scores and hierarchical F1-scores on proteins of different EC numbers grouped by the number of training samples available for their respective EC numbers. When an EC number has more than 100 training samples, ACCESS's predictive F1-score stabilizes at a consistently high level.

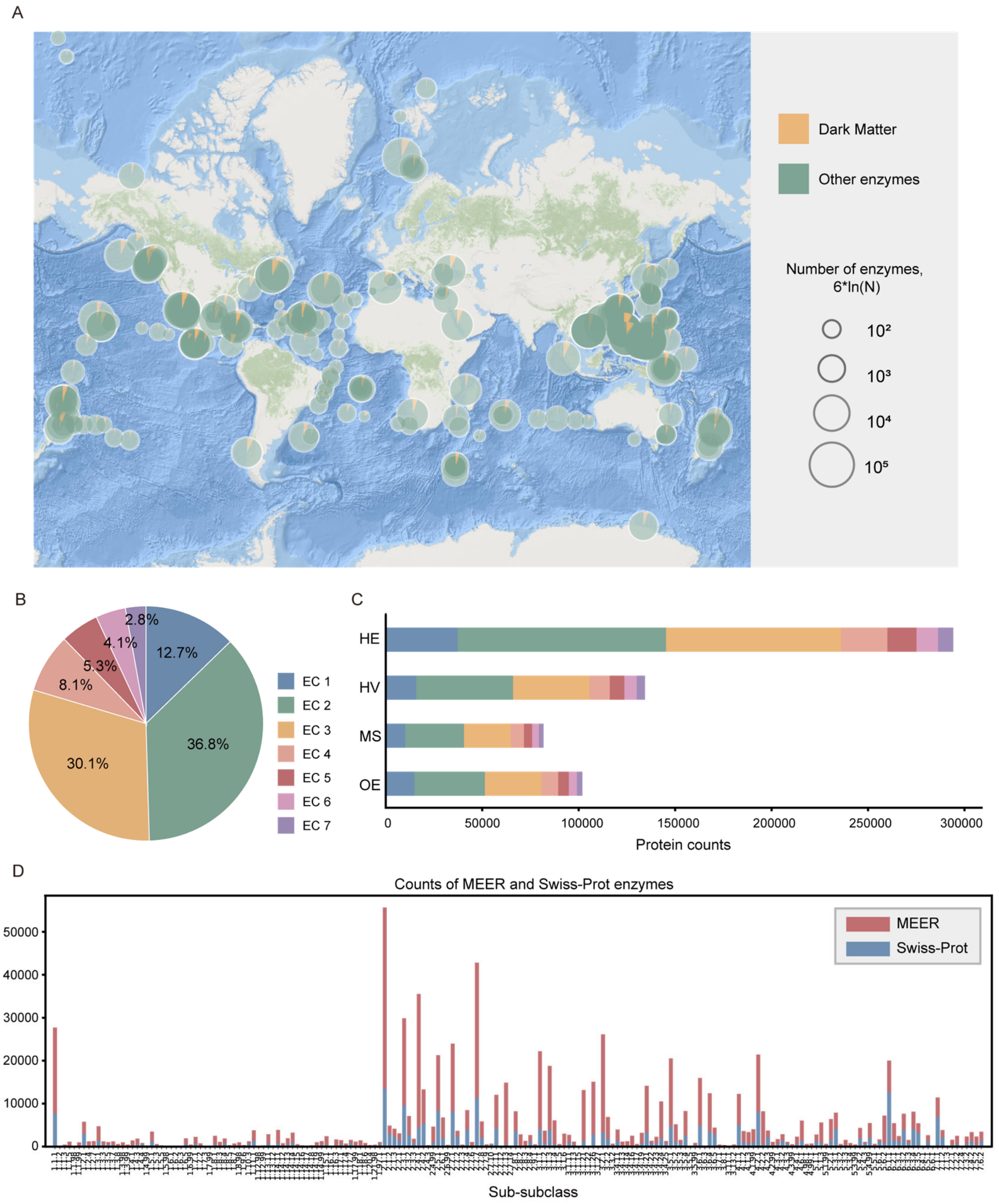


**Figure S3. Additional results on the atlas of enzymes from extreme environments, related to Figure 3.**

(A) Global biogeography of "functional dark matter" proportion. Overlaid pie charts on a world map illustrating the geographical distribution of "functional dark matter" enrichment across latitudes and longitudes, from the Arctic to the Antarctic, all major oceanic regions, and habitats ranging from shallow coastal areas to the Mariana Trench.

(B, C) Functional landscape of newly discovered deep-sea enzymes. (B) Pie chart summarizing the functional distribution of the 612,310 novel enzymes annotated by ACCESS from the MEER dataset. (C) Bar charts breaking down the EC-class counts across four distinct deep-sea habitats.

(D) Stacked-bar chart for every EC sub-subclass, showing the number of Swiss-Prot proteins (bottom blue segments) and the newly predicted MEER proteins (top red segments).

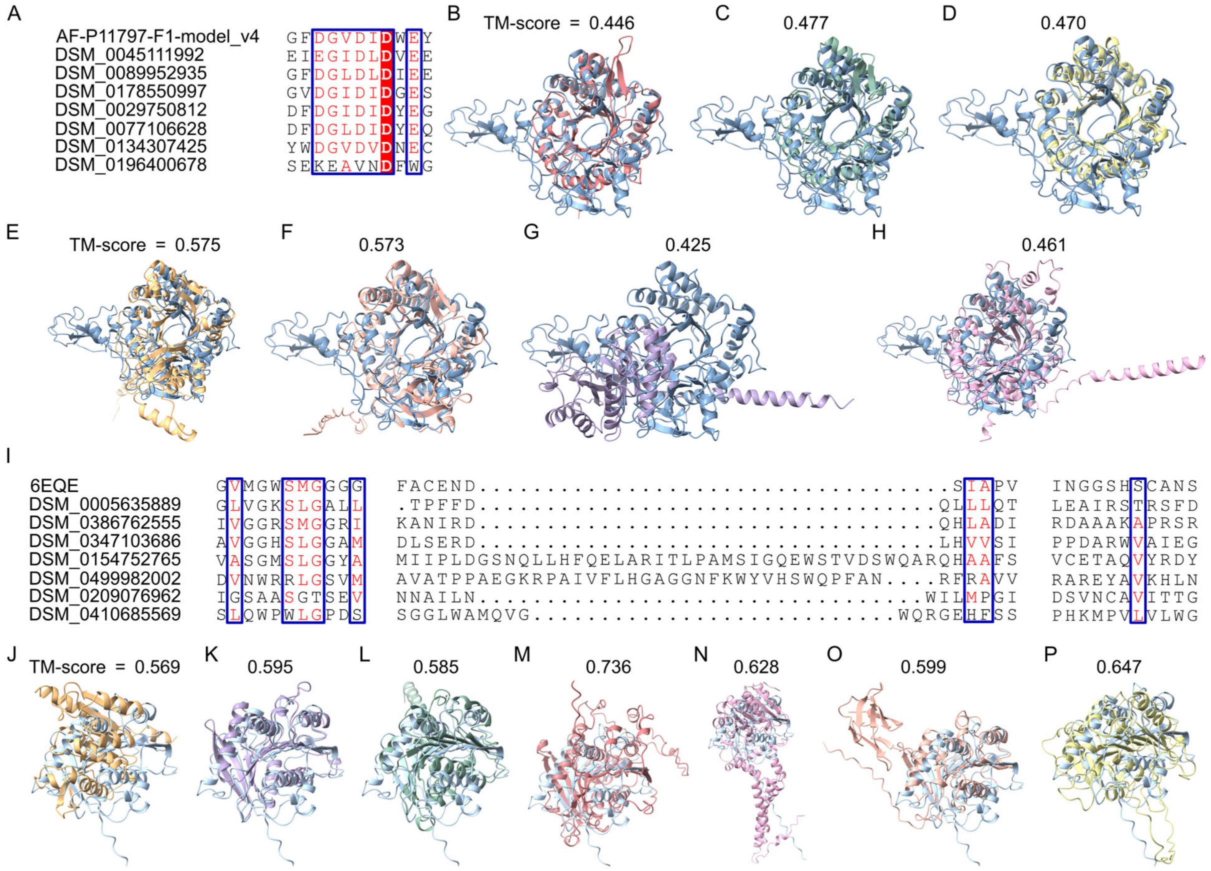


**Figure S4. Comparison of sequence and structure similarity between detected chitinase and PET hydrolase from MEER and the well-known proteins from the Swiss-Prot database, related to Figure 4.**

(A) Multiple sequence alignment of detected chitinases from MEER and the well-known chitinase from Swiss-Prot (AFDB ID: AF-P11797-F1-model_v4). Despite the difference in the overall sequences, the active sites are conserved.

(B)-(H) Overlap of chitinase structures detected from MEER and the well-known structure from Swiss-Prot and the corresponding TM-score normalized by the known structure. The overall structures are different while the region around active sites is still conserved.

(I) Multiple sequence alignment of detected PET hydrolases from MEER and the well-known PET hydrolase from Swiss-Prot (PDB ID: 6EQE).

(J)-(P) Overlap of PET hydrolase structures detected from MEER and the well-known structure from Swiss-Prot and the corresponding TM-score normalized by the known structure.

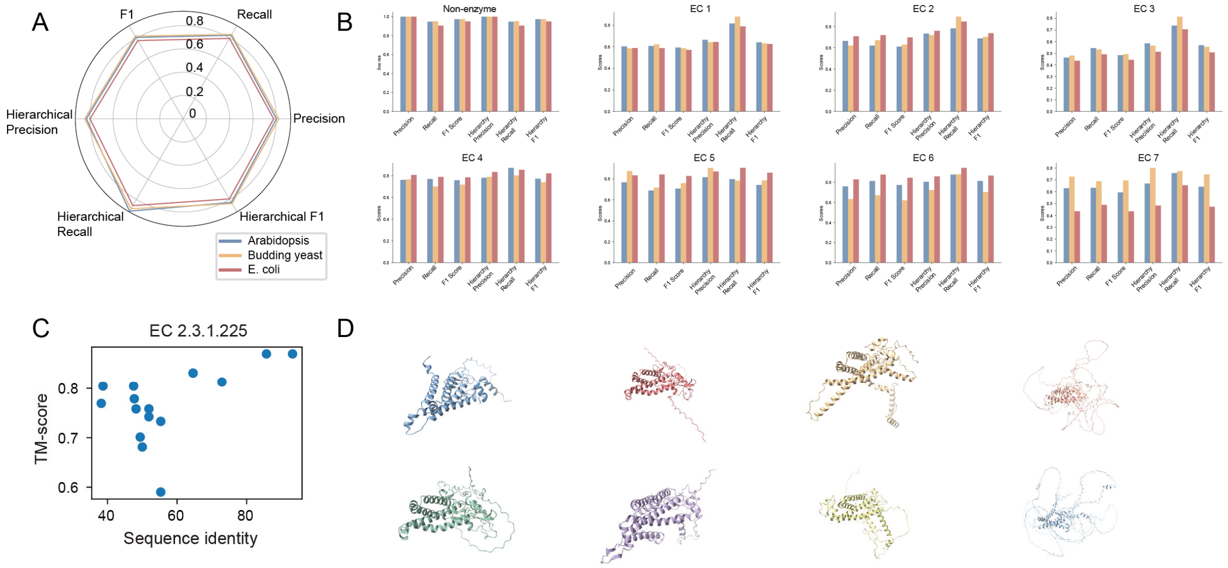


**Figure S5. ACCESS Reveals Macroevolutionary Trends and Functional Phylogenomics Across the Tree of Life, related to Figure 6.**

(A) A radar chart quantifying the consistency between functional predictions generated by ACCESS and annotations from the gold-standard resource, the Swiss-Prot database. The chart evaluates six key performance metrics for proteins from *Arabidopsis, Budding yeast* and *E. coli*, including precision, recall, F1-score, hierarchical precision accounting for EC hierarchy, hierarchical recall, and hierarchical F1-score.

(B) Performance metrics for non-enzymatic and all seven main EC classes of proteins from *Arabidopsis, Budding yeast* and *E. coli*, confirming the unbiased nature of ACCESS's high prediction accuracy.

(C) Distribution of 17 human proteins assigned to EC 2.3.1.225 (alpha-1,4-glucan branching enzyme) exhibiting divergence in sequence identity and TM-score (a measure of structural similarity).

(D) Visualization of 3D structural models of eight representative proteins from the EC 2.3.1.225 set in (C) with structural variation.
